# MRT-ModSeq – Rapid detection of RNA modifications with MarathonRT

**DOI:** 10.1101/2023.05.25.542276

**Authors:** Rafael de Cesaris Araujo Tavares, Gandhar Mahadeshwar, Han Wan, Anna Marie Pyle

**Author notes:** Equal contributions (Rafael de Cesaris Araujo Tavares, Gandhar Mahadeshwar, Han Wan).

## Abstract

Chemical modifications are essential regulatory elements that modulate the behavior and function of cellular RNAs. Despite recent advances in sequencing-based RNA modification mapping, methods combining accuracy and speed are still lacking. Here, we introduce MRT-ModSeq for rapid, simultaneous detection of multiple RNA modifications using MarathonRT. MRT-ModSeq employs distinct divalent cofactors to generate 2-D mutational profiles that are highly dependent on nucleotide identity and modification type. As a proof of concept, we use the MRT fingerprints of well-studied rRNAs to implement a general workflow for detecting RNA modifications. MRT-ModSeq rapidly detects positions of diverse modifications across a RNA transcript, enabling assignment of m1acp3Y, m1A, m3U, m7G and 2’-OMe locations through mutation-rate filtering and machine learning. m1A sites in sparsely modified targets, such as MALAT1 and PRUNE1 could also be detected. MRT-ModSeq can be trained on natural and synthetic transcripts to expedite detection of diverse RNA modification subtypes across targets of interest.

## INTRODUCTION

Post-transcriptional endogenous RNA modifications have been found to be integral to the robust functioning of many cellular processes including RNA stability (Boo and Kim, 2020; Kawata and Akimitsu, 2021; Wang et al., 2014), splicing (Karijolich et al., 2010; Martinez and Gilbert, 2018), mRNA export (Lesbirel et al., 2018; Roundtree et al., 2017b), translation (Eyler et al., 2019; Hoernes et al., 2016; Lin et al., 2016; Mao et al., 2019; Meyer et al., 2015), and protein metabolism and synthesis rates (Blanco et al., 2016; Guzzi et al., 2018). The landscape of such modifications is dynamic and tightly regulated so that the cell may adapt to the ever-changing nature of its microenvironment (Li and Mason, 2014; Licht and Jantsch, 2016; Roundtree et al., 2017a; Srinivas et al., 2021). In turn, the dysregulation of these modification patterns due to abnormal expression of RNA modifying enzymes has been linked to many developmental diseases, as well as the proliferation, growth, differentiation, and drug resistance of cancer cells (Barbieri and Kouzarides, 2020; Jonkhout et al., 2017; Nombela et al., 2021; Wood et al., 2021).

To study the function of RNA modifications and their regulatory networks, reliable and sensitive detection methods are desired. Although mass spectrometry technologies have the capability of detecting virtually all kinds of chemical modifications present in a sample, they lack the sensitivity and throughput necessary for the analysis of complex transcriptomic samples (Wetzel and Limbach, 2016). For this reason, the development of next-generation sequencing (NGS) approaches has gained importance for mapping of RNA modifications (Kumar and Mohapatra, 2021; Li et al., 2016). Several studies have employed high-throughput sequencing to detect specific types of RNA modifications, although the experimental protocols behind these methods range widely (Begik et al., 2021; Helm and Motorin, 2017; Leger et al., 2021; Motorin and Helm, 2019). Detection of pseudouridines (ψ), for example, is usually accomplished via functionalization of ψ sites as bulky CMC adducts to induce reverse transcription stops (Carlile et al., 2015). 2’-O-methylated nucleotides (Nm) lack a free 2’ hydroxyl group, and a method called Nm-seq (Dai et al., 2017) exploits this characteristic by utilizing periodate oxidation followed by phosphate elimination to enrich 3’ ends of transcript fragments at Nm sites prior to sequencing library construction. These methods require chemical treatment of the starting biological sample, they involve multistep protocols and many are stop-based methods, all of which limit throughput and diversification (Motorin and Marchand, 2021; Ryvkin et al., 2013). Most approaches are optimized for detection of a single type of RNA modification (Enroth et al., 2019; Li et al., 2017; Thalalla Gamage et al., 2021), which usually precludes their use for simultaneous profiling of multiple types of RNA modifications with the same experimental workflow.

Recently, the application of machine learning has facilitated detection of RNA modifications from RT signature data (Schmidt et al., 2019; Vandivier et al., 2019; Werner et al., 2020). Although these methods bypass the need for pre-treatment of the RNA sample, they have so far been limited to detection of only a few modifications on the Watson-Crick face, such as m1A and m1G. In addition, the low processivity of the RT enzymes employed in these methods makes them unsuitable for long RNA molecules.

The group II intron maturase of *Eubacterium rectale* (MarathonRT, or MRT) is an extremely processive reverse transcriptase enzyme (Zhao et al., 2018). It has been recently shown that MRT responds to the presence of certain chemical modifications on cellular RNAs by encoding mutations in the cDNA and that this behavior can be modulated using a non-natural divalent cofactor in order to selectively decrease reverse transcription fidelity at modified nucleotides (Guo et al., 2020). Here, we report MRT-ModSeq, a method for simultaneous positional and chemical assignment of diverse RNA modification subtypes that is applicable for long-read modification sequencing. We show that the preferential incorporation of cDNA mismatches by MRT under reaction conditions containing different divalent cations produces statistical patterns of mutation that are diagnostic of specific RNA modification subtypes. The signatures obtained with this methodology reveal the specific positions of common modified nucleotides within a transcript. With the training sets currently available, the actual chemical identity of many common modifications, such as m1acp3Y, m3U, m7G, m1A and m3U and Nm (2’-Ome modifications) can be specifically assigned. We used the annotated mutational fingerprints of well-characterized cellular RNAs as training features in a supervised machine learning pipeline. The strategy for data generation and machine learning implementation described in this work is adaptable and can be refined by incorporating new training signatures for identification of diverse RNA modifications in target transcripts.

## RESULTS

### A mutational profiling strategy for RNA modification detection with MRT

Given the reported ability of MRT to preferentially incorporate mutations in the cDNA at chemically modified nucleotides (Guo *et al*., 2020), we set out to generate and examine MRT mutational profiles for abundant cellular RNAs using a dual divalent cation approach devised in our laboratory. In this experiment, reverse transcription of target RNAs is performed separately in the presence of either magnesium or manganese and followed by library construction, producing an initial mutational profile in two separate dimensions, i.e., Mg and Mn mutation rates (Fig 1). As an initial approach, we used two highly abundant and well-studied transcripts as initial model systems: human 18S and 28S ribosomal RNAs, both of which harbor an abundance of 2’-O-methylated (Nm) and pseudouridine (ψ) sites, among other types of RNA modifications (Sloan et al., 2017; Taoka et al., 2018). We were able to obtain highly reproducible mutational patterns across independent experiments (Figs S1-S2) in a variety of human cell lines for both rRNAs (Figs S3-S6), suggesting that MRT mutation rates of individual nucleotide sites are highly replicable for both Mg and Mn conditions.

**Figure 1.**
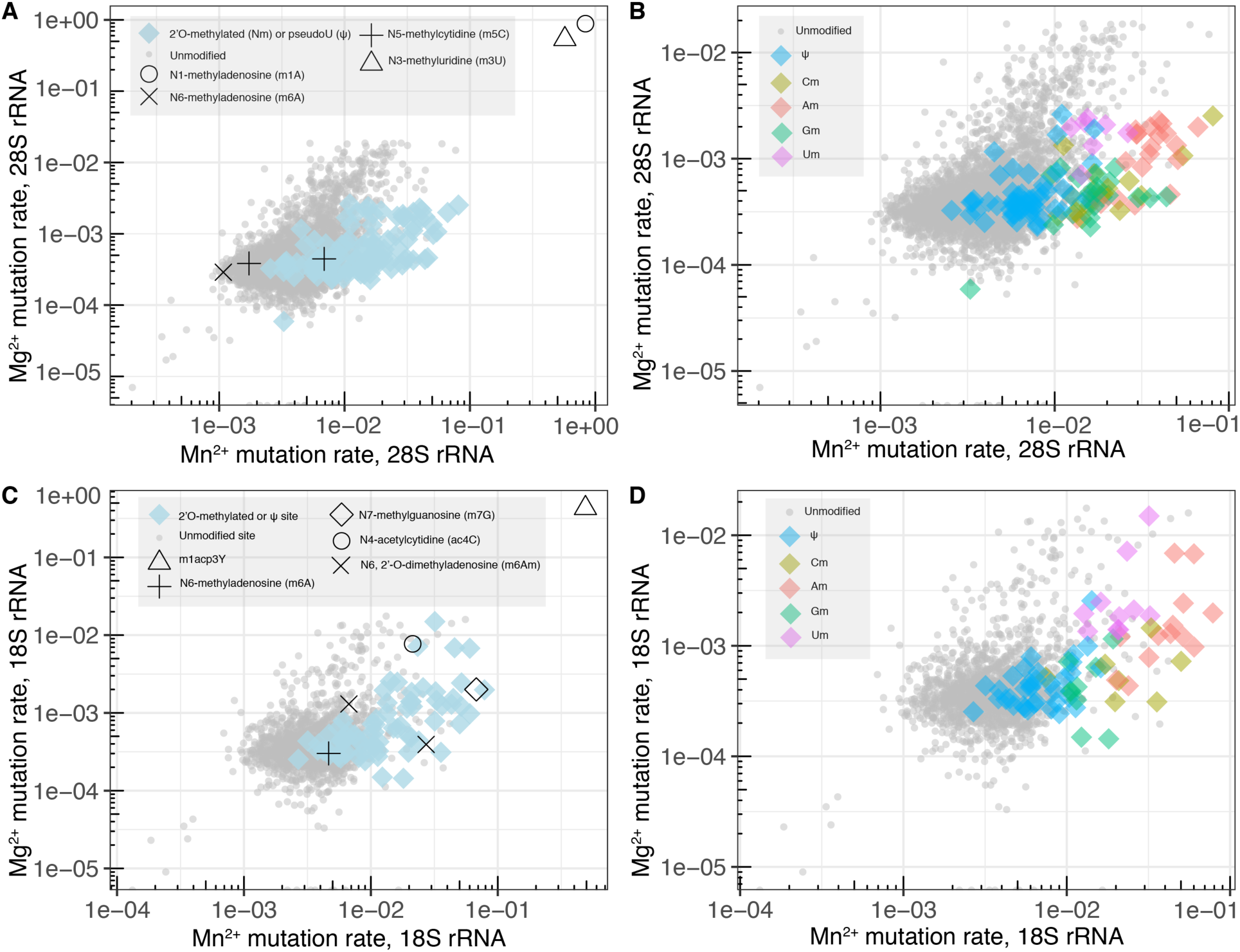
2-D mutational profiles obtained with MRT in the presence of Mg or Mn for human 18S and 28S rRNAs. (A) Log-scale plot of Mg (y-axis) and Mn (x-axis) mutation rates obtained with MRT for the human 28S rRNA. Each group of nucleotides (unmodified and different types of modified) are represented by different symbols as indicated in the legend. (B) Version of (A) plot including only Nm and ψ modifications, in addition to unmodified sites. Different types of Nm are indicated by different colors, according to the legend. (C) Log-scale plot of Mg (y-axis) and Mn (x-axis) mutation rates obtained with MRT for the human 18S rRNA. Each group of nucleotides (unmodified and different types of modified) are represented by different symbols as indicated in the left-hand legend. (D) Version of (C) plot including only Nm and ψ modifications, in addition to unmodified sites. Different types of Nm are indicated by different colors, according to the legend.

On average, Mn mutation rates are greater than Mg mutation rates by almost one order of magnitude, i.e., MRT is generally more susceptible to incorporating mutations in the presence of Mn than in the presence of Mg (Fig 1). Importantly, this behavior is exacerbated in the case of nucleotides harboring RNA modifications, i.e., overall, modified sites exhibit greater Mg-Mn mutation rate differences than unmodified sites. As a result, we observed varying degrees of mutational segregation between modified and unmodified nucleotides, and the amount of segregation largely depends on the identity of the chemical modification. When examining the signatures of human 18S and 28S rRNAs (Figs 1A, 1C), we were able to categorize three main groups of modified nucleotides based on their mutational segregation relative to unmodified sites: sites with average mutation rates roughly two orders of magnitude greater than those of other sites, which included three base methylations, m1acp3Y, m3U, and m1A; sites with average mutation rates almost one order of magnitude greater than other sites, including one base methylation (m7G), one base acetylation (ac4C) and the ribose 2’-OH methylations (Nm); and sites with average mutation rates essentially indistinguishable from other sites, which included ψ (pseudouridine), and three other base methylations (m5C, m6A and m6Am). Although most types of RNA modifications in our initial 18S/28S transcript set are present at single or very few instances, 2’-O-methylations (Nm) and pseudouridines (ψ) are found in multiple positions, which allowed for a more detailed group analysis (Figs 1B, 1D). Interestingly, we observed that different nucleotides bearing 2’-OH methylations (Am, Gm, Cm, Um) produce different mutational signatures (Figs 1B, 1D), which not only suggests that Nm sites effectively behave as four distinct types of modifications in this experiment, but also indicates that these sites should be categorized into four separate classes in any downstream analysis of their mutational profiles.

### MRT mutation rates are highly dependent on modified nucleotide identity

To better understand the mutation rate distributions produced by MRT in response to the most abundant rRNA modifications, Nm and ψ, we examined the 2-D mutational profiles (Mg, Mn) of these sites when classified by nucleotide identity. As anticipated, we found that each type of nucleotide occupies a different region in the mutational space defined by these two variables (Fig 2). Unmodified sites of each nucleotide type exhibit generally similar behavior: the majority of unmodified nucleotides have Mg^2+^ mutation rates between 1e-04 and 1e-03, and Mn^2+^ mutation rates between 1e-03 and 1e-02, consistent with the basal increase in positional mutation rate in the presence of Mn^2+^. Interestingly, clusters of unmodified nucleotides exhibit a sparse tail of sites with higher mutation rate values, where the Mn^2+^ mutation rate is approximately commensurate with the Mg^2+^ mutation rate, a pattern that roughly defines a ‘mutational trajectory’ that is common to all types of unmodified nucleotides. Similarly, each group of Nm sites forms a distinct ‘trajectory’ in this 2-D space. In the case of Am sites, we see minimal overlap between modified and unmodified dispersions, which results in the highest mutation segregation among Nm groups. This is mostly due to a pronounced shift in the Mn dimension (Fig 2A). Cm and Gm distributions show reduced spreads in one of the two dimensions when compared to their unmodified counterparts, resulting in spatial distributions far less parallel than in the Am case. Specifically, Cm sites show lower spread in the Mg axis (Fig 2C), while Gm sites show lower spread in the Mn axis (Fig 2D) relative to their respective unmodified distributions. Curiously, the ‘trajectories’ described by Um sites and unmodified U sites appear to largely fall the same line, i.e., the two distributions belong to the same apparent (diagonal) trend, which is unique to this group (Fig 2B). In contrast with Um nucleotides, pseudouridines (ψ) occupy a region in mutational space that largely overlaps with that of unmodified uridines, indicating that no deconvolution is readily achievable in these two dimensions under the experimental conditions we employed.

**Figure 2.**
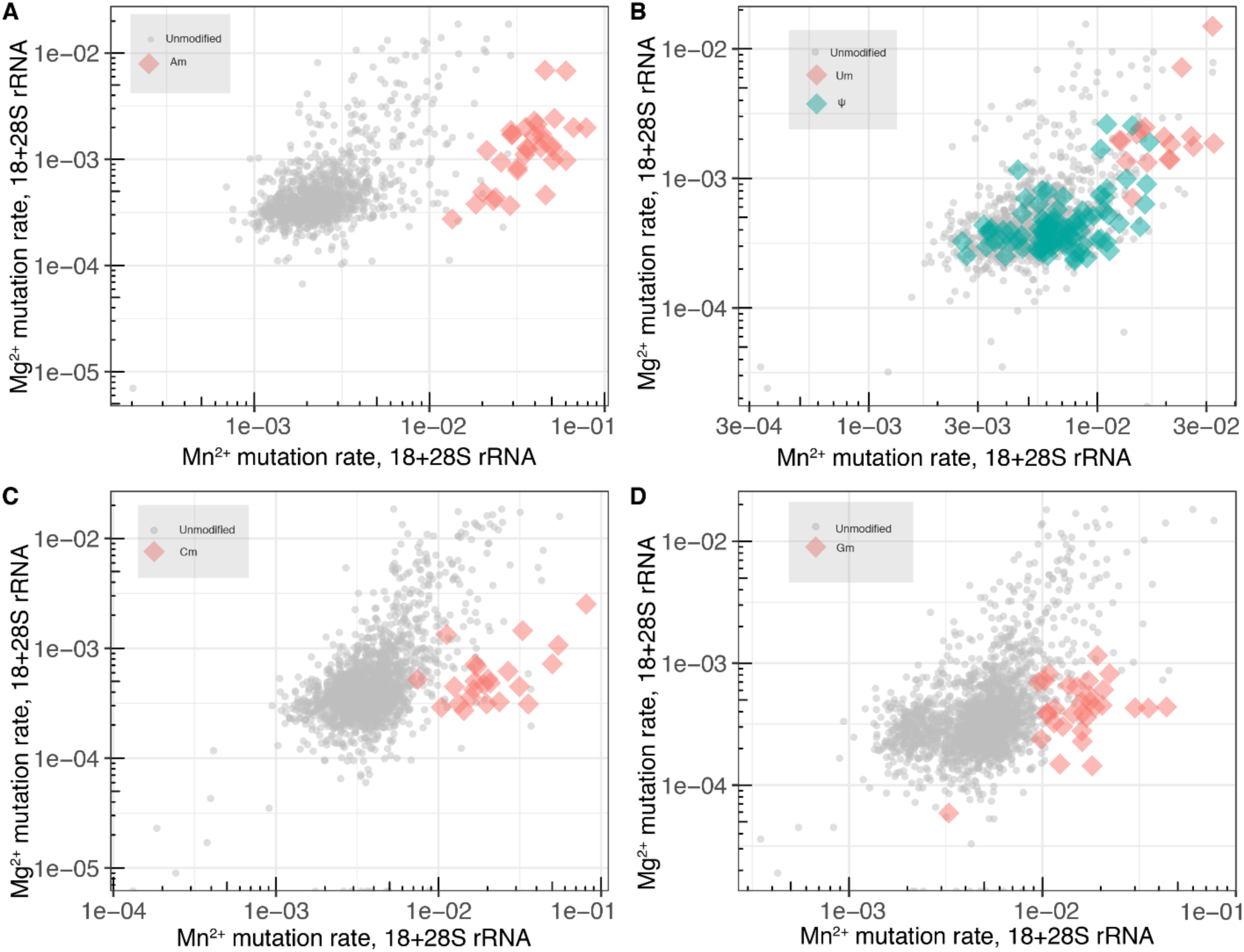
Combined 2-D mutational profiles obtained with MRT in the presence of Mg or Mn showing data for both 18S and 28S rRNAs. (A) Log-scale plot of Mg (y-axis) and Mn (x-axis) mutation rates obtained with MRT for the combined 18S/28S dataset showing Am (orange) and unmodified A (grey) sites, only. (B) Log-scale plot of Mg (y-axis) and Mn (x-axis) mutation rates obtained with MRT for the combined 18S/28S dataset showing Um (orange), ψ (green) and unmodified U (grey) sites, only. (C) Log-scale plot of Mg (y-axis) and Mn (x-axis) mutation rates obtained with MRT for the combined 18S/28S dataset showing Cm (orange) and unmodified C (grey) sites, only. (D) Log-scale plot of Mg (y-axis) and Mn (x-axis) mutation rates obtained with MRT for the combined 18S/28S dataset showing Gm (orange) and unmodified G (grey) sites, only.

### Mutation compositions significantly differ between modified and unmodified sites

Knowing that modified sites differ in their mutation patterns from unmodified sites (Fig 1) in a nucleotide-dependent manner (Fig 2), we proceeded to dissect MRT cDNA mismatch signatures in order to gain insights into how the enzyme responds to the presence of different modifications in the RNA template. As in the previous case, we focused this analysis on the abundant Nm and ψ modifications present in our rRNA dataset. First, we computed marginal mutation rates for both modified and unmodified sites by calculating mismatch rate compositions sorted by nucleotide type (e.g., A-to-C mut rates, A-to-G mut rates, etc). These marginal mutation rates were then scaled and normalized by the overall mismatch rate to yield a series of “mutational fingerprints” (Fig 3) and facilitate visualization of the relative contributions of each type of mismatch event to the overall mismatch pattern.

**Figure 3.**
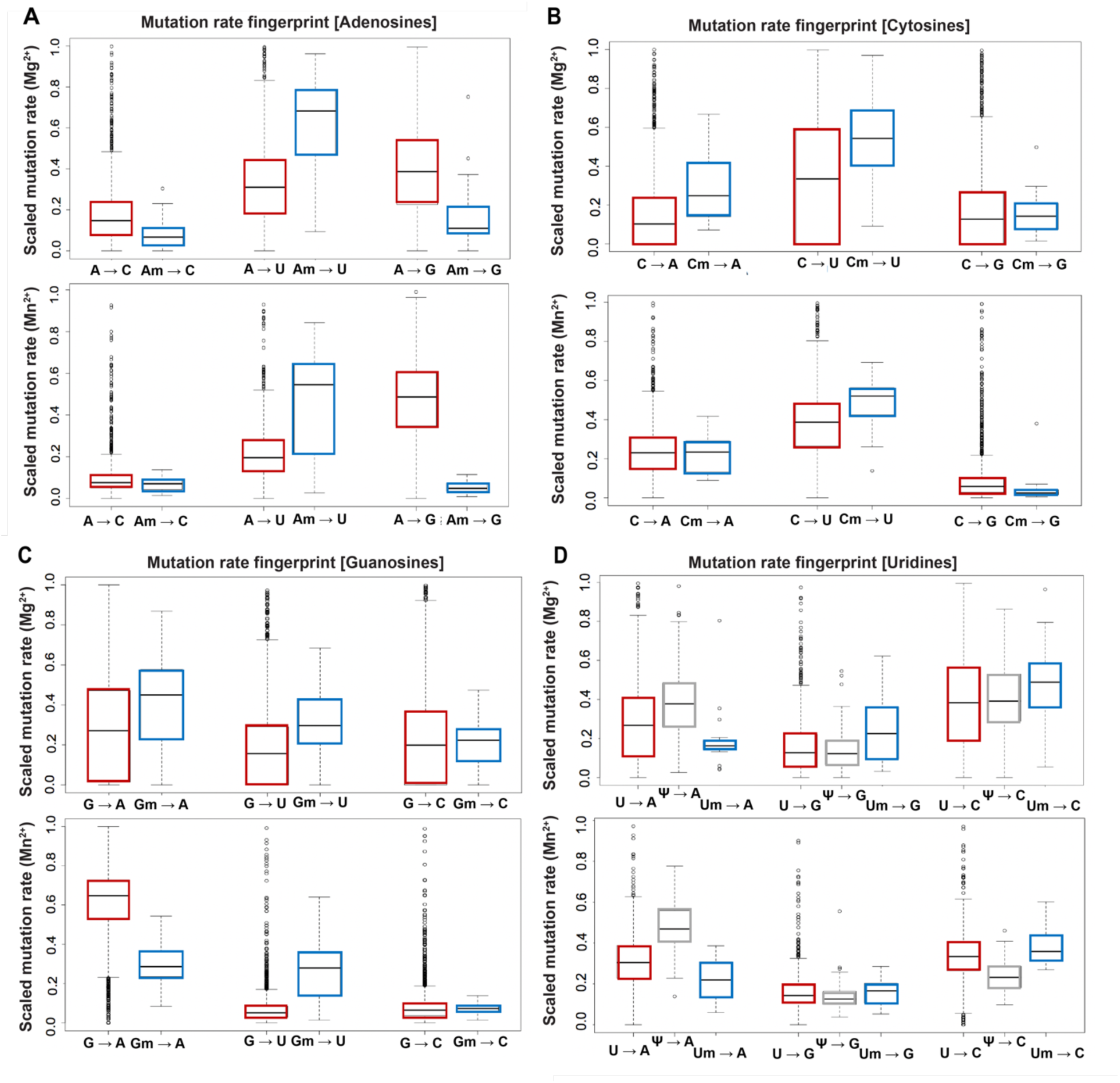
Marginal mutation rate analysis of the combined 18S/28S rRNA dataset. (A) Box and whiskers plot of marginal (scaled) mutation rates calculated from the Mg sample (upper graph) and Mn sample (lower graph) for adenosine sites. Unmodified A sites are represented in red and Am sites are in blue. (B) Box and whiskers plot of marginal (scaled) mutation rates calculated from the Mg sample (upper graph) and Mn sample (lower graph) for cytosine sites. Unmodified C sites are represented in red and Cm sites are in blue. (C) Box and whiskers plot of marginal (scaled) mutation rates calculated from the Mg sample (upper graph) and Mn sample (lower graph) for guanosine sites. Unmodified G sites are represented in red and Gm sites are in blue. (D) Box and whiskers plot of marginal (scaled) mutation rates calculated from the Mg sample (upper graph) and Mn sample (lower graph) for uridine sites. Unmodified U sites are represented in red, pU (ψ) sites are in grey and Um sites are in blue.

Overall, we noticed that most marginal mutation rates were either elevated or unchanged relative to unmodified sites, although we also found a few instances of decreased marginal mutation rates (Fig 3). Remarkably, we observed the biggest differences between Nm and unmodified sites at purine sites (Am, Gm), while pyrimidine nucleotides (Cm, Um) exhibited more modest changes (Fig 3), suggesting that the additional bulkiness imparted by a 2’-O-methyl group has a bigger effect on MRT reverse transcription fidelity in the case of purine sites than in the case of pyrimidine sites. Upon closer inspection of the signatures of 2’-O-methylated purines, we identified a common pattern for As and Gs. At Gm sites, we detected considerably higher G-to-U mismatch rates in the presence of Mn^2+^ along with decreased G-to-A mismatch rates (Fig 3C), relative to unmodified Gs; on the cDNA level, these correspond, respectively, to enhanced misincorporation of As and decreased misincorporation of Ts (pyrimidine). At Am sites, we observed significantly elevated A-to-U mismatch rates in the presence of either Mg^2+^ or Mn^2+^, along with decreased A-to-G mismatch rates, relative to unmodified As (Fig 3A); in this case, these changes correspond to increased misincorporation of As and decreased misincorporation of Cs (pyrimidine), respectively. The relative trends in each case are seemingly opposite (e.g., A-U goes up, while A-G goes down), since all marginal mutation rates (relative contributions to the total) add up to 1. We conclude that the MRT signature of purine Nm sites (Am, Gm) consists of a specific preference towards A-misincorporation and a concomitant decrease in pyrimidine-misincorporation on the cDNA strand.

Although pyrimidine sites (Cs and Us) exhibited less pronounced signature differences between Nm and unmodified nucleotides, we did observe slightly elevated C-to-A (Mg) and C-to-U (Mg, Mn) rates at Cm sites (Fig 3B). In contrast, marginal mutation rates for Um sites were almost unaffected compared to unmodified sites (Fig 3D), suggesting that even though there is a global increase in the overall mutation rates, the Um mutational composition is largely similar to that of unmodified Us. Distinct from purine sites, we did not see a particular preference towards either purine or pyrimidine cDNA misincorporation for either Cm or Um.

While it was not possible to glean significant mutational segregation for pseudouridines relative to unmodified uridines based in a 2-D mutational space (Fig 2B), upon examination of marginal mutational signatures we observed that pseudouridines exhibit elevated U-to-A mismatch rates in the presence of Mn^2+^ and depressed U-to-C rates (Fig 3D). In this case, there was no correlation between the observed misincorporation preferences and nucleobase bulkiness, consistent with the similar sizes of uridines and pseudouridines, but these trends suggest that pseudouridines fingerprints might still be useful for detection purposes.

### Nm and ψ fingerprints are largely independent of sequence context

To assess the context dependence of mutational fingerprints and evaluate potential sequence biases in MRT-derived signatures of Nm and ψ sites, we categorized the rRNA dataset using sequence-based criteria. For this purpose, we denote the ‘penultimate nucleotide’ as the nucleotide immediately 3’ to the nucleotide under consideration. We then sorted marginal mutation rates for each modified nucleotide according to its penultimate nucleotide identity (Fig 4, Fig S7). For each group of modified nucleotides, Mn mutational signatures are remarkably conserved regardless of the penultimate nucleotide identity (Fig 4), suggesting a relatively minor influence of the underlying sequence context on mutational composition. Similarly, Mg mutational patterns were, in most cases, only very slightly affected by penultimate nucleotide identity (Fig S7), implying that Nm and ψ fingerprints show minimal variation as a function of their immediate downstream sequence context. Additionally, we looked at the influence of the post-terminal nucleotide, i.e., the nucleotide immediately upstream of the nucleotide under consideration, in both Mg and Mn cases (Figs S8-9) and confirmed the absence of significant biases towards neighboring nucleotide types in that case. These observations indicate that fingerprints obtained at those specific sites directly report on their chemical modification status and are not significantly dependent on downstream or upstream nucleotide identity. Given the large amount of information encoded in the RT fingerprints obtained through this strategy, we surmised that we might be able to utilize the differential mutation signatures of modified sites as indicators of the presence of specific RNA modifications in a classification framework.

**Figure 4.**
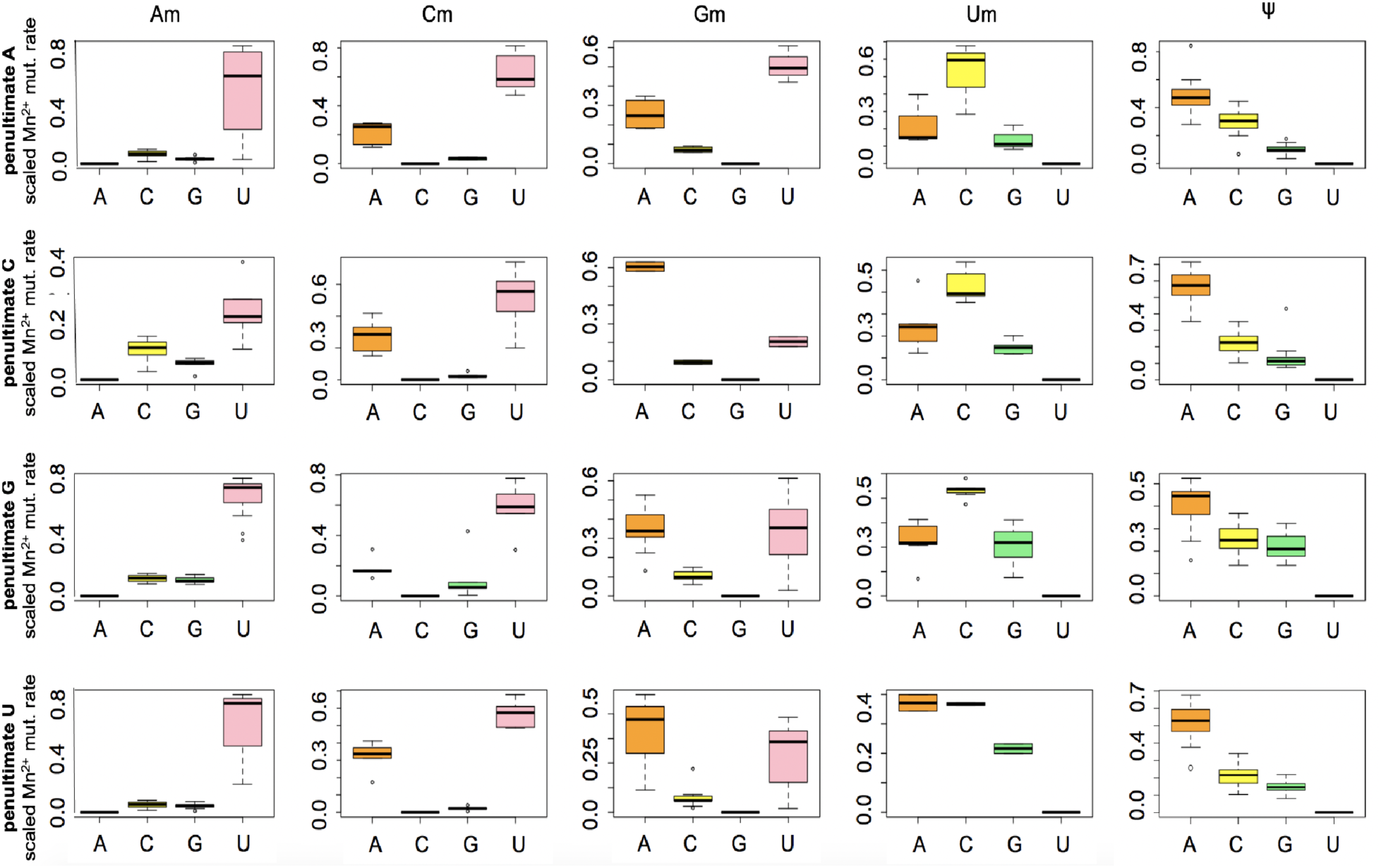
Sequence context analysis of manganese (Mn) mutational signatures of Nm and ψ sites in the combined 18S/28S rRNA dataset. The results are sorted by modified nucleotide type (columns) and by the penultimate nucleotide in the RNA sequence, i.e., the nucleotide immediately 3’ to the position being examined. Each individual plot shows the distribution of marginal (scaled) mutation rates calculated from the Mn sample as a function of the different groups of mutated nucleotides (e.g., the top graph in the Am column shows marginal mutation rates from A to each one of the 4 nucleotides: A to A (no mutation), A to C, A to G, A to U).

### MRT-ModSeq dataset preparation and training workflow

In order to refine and improve our ability to predict RNA modifications on a target RNA, we used MRT-derived mutation rates from both Mg and Mn reactions to implement a machine learning framework (Fig 5). Since we could readily obtain high-depth sequencing data using MRT for both human 18S and 28S rRNAs, these RNAs were used to build an initial class training set to classify RNA modifications within this method (Fig 5 – steps 1-3). After scanning mutation rates in our training set, we found certain types of base modification (m1acp3Y, m3U, and m1A) to be very highly mutagenic (Mn mutation rate > 50%, see Fig 1). Because only a handful of these sites exist in our training set, we cannot use these for robust classification in the machine learning setting, but we can scan any test RNA transcript using a high-pass filter for statistically significant levels of misincorporation that are consistent with the presence of m1acp3Y, m3U, or m1A modifications. In addition to these, we included a filter for potential m7G sites (see Methods), since these also exhibit mutational signatures that are significantly different from other G sites (unmodified and Gm) present in the training set (Fig 1). Once these sites are detected and filtered out of the set, downstream analysis consists of using machine learning to classify the remaining RNA modifications. As a proof of concept, the human 18S/28S rRNA training set enabled an ML implementation for classification of Nm and ψ sites, since these modifications were present in multiple instances throughout both RNAs. Here, we describe this implementation in detail.

**Figure 5.**
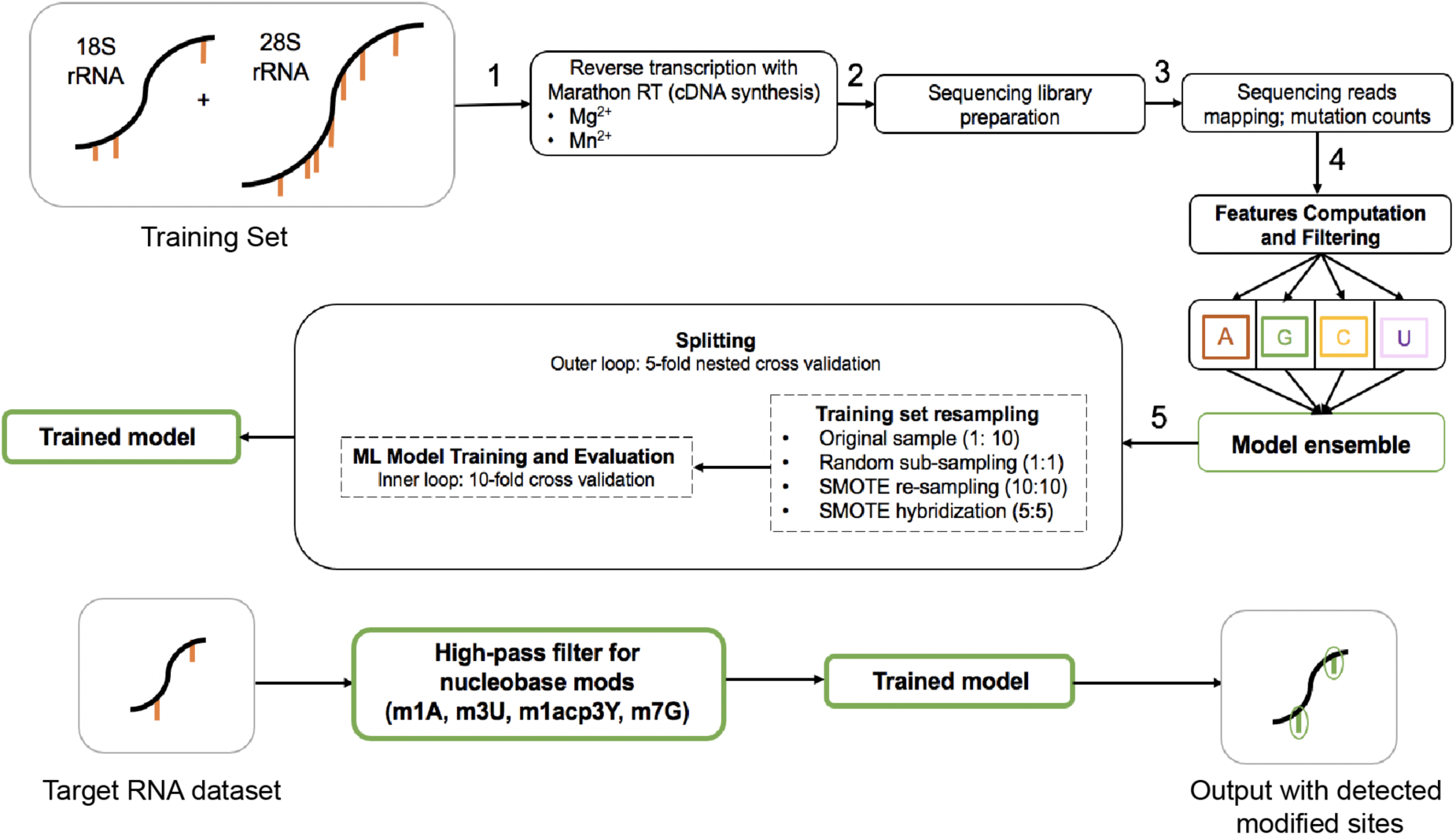
MRT-ModSeq workflow. The scheme illustrates the pipeline using human 18S and 28S rRNAs to obtain training datasets by (step 1) carrying out the RT reaction separately in the presence of either Mg or Mn. The resulting cDNA is used to (step 2) prepare sequencing libraries; the sequencing reads obtained are used to count mutations at each nucleotide position (step 3), which is followed by filtering out poorly covered positions and computing nucleotide changes (features) and partitioning the dataset into four different groups that compose the model ensemble (step 4). Training (step 5) is carried out by benchmarking different ML algorithms in a nested cross-validation routine where the best performing algorithms are chosen for each class to generate the final trained model (step 6). The diagram in the bottom shows the MRT-ModSeq workflow for detection of modified sites in a target RNA.

Since mutation signatures are unique to nucleotide type in the case of Nm modifications, we reasoned that Nm detection must occur through four distinct classification algorithms, each trained on and used to classify Nm sites of the same nucleotide type. Additionally, given that marginal mutation rates represent fingerprints of distinct classes of modified nucleotides (Figs 3-4), and are therefore variables that might increase classificatory power, they were also included as predictors in the ML models in addition to overall mutation rates (see Methods). In order to correct for the severe imbalance of the training set (>95% of the sites in each dataset are unmodified), we also tested different methods of synthetic minority oversampling (SMOTE) (Blagus and Lusa, 2013; Chawla et al., 2002). After splitting our original dataset in a 5-fold cross-validation schema to prevent data leakage from synthetic samples, the outer loop training set was resampled in three different ways (Fig 5 – step 5): an undersampled test, where the majority class (unmodified sites) is randomly subsampled until its equally weighted with the minority class; an oversampling test, where the minority class was resampled to match the weight of the majority class; and a hybrid test, utilizing both oversampling of the minority class and subsampling of the majority class to equate class weights. Using this strategy, we benchmarked several ML algorithms (see Methods) along with bagged and boosted versions of each algorithm (15 algorithms in total) on each training set in the resampling loop using an inner 10-fold cross validation loop. This process was repeated for each nucleotide type and performance was scored in each case. After benchmarking, AdaBoost paired with tree-based methods resulted in the best performance for adenosines and cytidines, while bagged tree-based methods were the best performers on guanosines and uridines. Importantly, SMOTE did not uniformly improve classificatory power across all nucleotide groups (Tables S8-S11). In some cases, solely re-sampling the minority class using SMOTE tended to overfit the data and negatively affected performance. This could be due to the large imbalance between modified and unmodified classes, since imputing synthetic samples to scale the minority class size by ∼100x might artificially bias the algorithm to learn certain synthetic characteristics not represented in the empirical sample. In turn, either the original dataset (e.g., adenosines, uridines, guanosines) or a hybridized dataset (e.g., cytidines) was used to train each prong of the algorithm, according to benchmarking performance.

Since different target RNAs might exhibit modification landscapes vastly different from those of the rRNAs that we trained our algorithms on and our training set predominantly consists of Nm and ψ modifications, we cannot readily extend an algorithm trained on these sites to RNA sequences harboring a different set of modifications. With this problem in mind, we designed a two-pronged algorithm implementation strategy, corresponding to two separate real-life scenarios. For the case in which we know that the test RNA transcript harbors mostly Nm and/or ψ, similar to the transcripts in our initial training set, we aimed for an algorithm that maximizes *sensitivity* (maximizes proportion of ‘positives’ that are correctly identified). We call this the permissive algorithm, as it attempts to correctly classify as many modified sites as possible. In the alternate use-case, where we do not know much about the modification landscape of the target transcript, or where we know that the transcript does not predominantly have Nm and ψ modifications, we wish to use an algorithm that maximizes specificity (maximizes proportion of ‘negatives’, or unmodified sites, that are correctly distinguished from modified sites). We called this the stringent algorithm, since it uses a cost matrix to impart greater penalty on false positives, and thus requires a higher burden of proof to classify sites as ‘modified’. Sites deemed modified by the stringent algorithm exhibit signatures that are consistent with the presence of RNA modifications similar or different from the ones used in the training set. The resulting MRT-ModSeq pipeline applies a high-pass filter to screen for certain nucleobase modifications (m1acp3Y, m1A, m3U, m7G), followed by an initial 18S/28S-trained model to yield candidate positions on target RNAs for Nm and ψ modifications.

### MRT-ModSeq benchmarking tests

To evaluate the performance of MRT-ModSeq, we conducted a routine of benchmarking tests. Initially, we assessed how ML models trained on one RNA containing a set of modifications would perform when used to detect similar modifications on a different transcript. This was also important for evaluating the consistency of mutation signatures across different RNAs for the same class of modifications. For this purpose, we independently generated a training set that included only 28S rRNA data to predict the modification sites on 18S rRNA (Table S1), and vice versa (Table S2). These “inter-set” tests were performed using sequencing data from independent experiments for the training and test RNAs to further simulate a “true” test case, where the dataset used to train the ML algorithm might be generated independently from the test RNA dataset. The first step in the MRT-ModSeq pipeline is to identify nucleobase modifications based on the mutational signatures of the test dataset. In inter-set tests, however, this cannot be tested as the highly mutagenic nucleobase modifications (m1acp3Y, m3U, m1A, m7G) are not commonly shared between 18S and 28S rRNAs. Rather, we tested how the ML algorithm performed when used to classify Nm and ψ sites (28S-trained vs 18S-test and 18S-trained vs 28S-test). Since, in this case, the same classes of modifications (Nm and ψ) were present in both training and test RNAs, we chose to use ModSeq’s permissive ML algorithm, as opposed to the case where no prior knowledge on the modification status of the test RNA is available, for which the stringent model would be more appropriate. Despite the heavily class-imbalanced nature of both training and test RNA sequences (number of unmodified >> number of modified sites), the algorithm accurately and precisely identified >99% of the sites as unmodified (low false discovery rate) and was selective enough to capture the majority of Nm sites in each target RNA (MCC ranging from 0.571 to 0.959, depending on nucleotide type, with the best performance observed for Am and Um classes). In contrast, the algorithm performance on ψ detection was rather limited (MCC < 0.4) in both cases (Tables S1-S2), consistent with the much lower cluster segregation between unmodified and pseudouridine sites in the training set (Fig 2).

After testing MRT-ModSeq in this more limited cross-set fashion, we next evaluated how it performed when used to detect modifications on RNAs other than the ones used as training set. To achieve this, we tested the complete pipeline on a series of abundant RNAs with experimentally verified modification landscapes (Noeske et al., 2015; Taoka *et al*., 2018; Yang et al., 2016): human 5.8S rRNA (Table S3); *E coli.*16S (Table S4) and 23S rRNAs (Table S5); and yeast 18S (Table S6) and 25S rRNA (Table S7). In these tests, MRT-ModSeq was trained on the combined human 18S/28S dataset and included both the high-pass filter for base modifications and machine learning steps.

The first group of RNAs consists of transcripts that are only sparsely modified. The first example, human 5.8S rRNA, has two positions of 2’-O-methylation, 14 (Um) and 75 (Gm), and two pseudouridines (positions 55 and 69). MRT-ModSeq mutation rates for this transcript were not consistent with the presence of m1A, m7G, m3U modifications, as expected from the absence of those in this RNA. Using the ML algorithm, the method accurately classified the two Nm sites of this transcript (Table S3), although the two pseudouridines were incorrectly classified as unmodified (false positives). The second example, *E. coli* 16S rRNA, is an interesting test case, since this RNA contains no Nm modifications and only 1 pseudouridine, in addition to a few base modifications. MRT-ModSeq successfully detected both m7G and m3U positions in 16S rRNA based on the high-pass mutation rate filter. Encouragingly, the ML algorithm only detected 3 false positives on the transcript - 1 false Gm and 2 false Um sites. Interestingly, the G site, identified on position 966, is in fact an m2G site, a base modification not present in our training set, which suggests that sites flagged by this algorithm might actually contain modifications whose fingerprints overlap with those of Nm sites and are thus suitable candidates for posterior validation. Also in this case, the single pseudouridine present in 16S rRNA was not detected by the ML algorithm (false negative). The last test RNA in this group, *E. coli* 23S rRNA, has only one m7G site, which was not detected by the high-pass filter of ModSeq. However, the filter identified two other candidate sites, one that is actually an m1G site and another with no known modification. All three Nm sites (positions 2251, 2499, and 2552) in 23S rRNA were accurately classified by the ML algorithm (Table S5). In addition, only two unmodified sites (U) were incorrectly classified as Nm (false positives). Finally, out of ten pseudouridine sites on the 23S rRNA, 3 were correctly identified (23S rRNA positions 1911, 1915, 1917). Importantly, in all three RNAs tested in this group, no Nm false negatives were observed, suggesting that the method operates with good sensitivity even in cases where the signal is scarce and sparsely distributed.

For the second group of test RNAs, *S. cerevisiae* 18S and 25S rRNAs were used since they are considerably more abundantly modified with Nm and ψ than the previous group. Yeast 18S rRNA in yeast harbors two of the base modifications we tested for - m7G and m1acp3Y - and both were detected by ModSeq mutation rate filter (Table S6). Out of its 8 Am positions, 7 were correctly identified by the algorithm (with only one false negative at position 796); out of 3 Cm sites, 2 were correctly identified (with one false negative at position 1007); out of 5 Gm sites, 3 were correctly identified (with two false negatives at positions 1428 and 562); finally, the algorithm did not catch the 2 Um positions in this RNA (false negatives). The second RNA in this group, yeast 25S rRNA, contains two m1A and two m3U sites, all of which were detected by our high-pass filters (Table S7). In the Nm case, 8 out of 12 Am nucleotides were correctly identified by the algorithm (with four false negatives at positions 649, 807, 1449, and 2280). Here, we found an interesting pattern with the Am sites missed by the algorithm: in all 4 cases of Am false negatives, there was another modified Nm site either immediately adjacent to the true Am or within a 2-nucleotide distance. It is possible that the presence of proximal or adjacent Nm sites influences MRT signatures. Accordingly, in each instance of Am with an immediately adjacent Nm, the adjacent Nm site was also not identified correctly, indicating that the local modification context might affect classification performance. 25S rRNA also harbors 7 Cm sites, 2 of which were correctly identified (with five false negatives at positions 650, 663, 2197, 2337, and 2959); 10 Gm sites, 5 of which were correctly identified (with five false negatives at positions 867, 1450, 2791, 2793, 2815); 8 Um sites, 5 of which were correctly identified (with three false negatives at positions 2347, 2421, 2921). Interestingly, one of these false negative Um sites (U2921) is adjacent to both a Gm and a pseudouridine, both of which were correctly identified. When we compare Nm classification results both groups of RNAs, the more highly modified group (yeast 18S and 25S rRNAs) returned fewer false positives and more false negatives than in the lightly modified group (human 5.8S, E coli 16S and 23S rRNAs). This trend is consistent with the fact that the presence in the second group of multiple shortly spaced or even consecutive modified nucleotides, which might affect detection sensitivity by reducing RT readthrough in densely modified clusters. In contrast with the overall good performance of Nm algorithms, pseudouridine detection showed comparably limited results in both groups of test RNAs, with an average MCC of 0.28 (Tables S3-S7, Fig S10). Given the largely overlapping signatures between ψ and unmodified U sites (Figs 1 & 2), the overall poor performance of ψ algorithms is most likely a result of the low degree of modified-unmodified segregation in both Mg and Mn RT conditions.

In addition to the highly abundant rRNAs used for our benchmarks, we also tested whether MRT-ModSeq could accurately detect RNA modifications in cases of sparsely modified, less abundant RNAs (e.g., mRNAs, lncRNAs). For this purpose, we selected two test RNAs containing experimentally-studied m1A modification sites: 1 – the MALAT1 lncRNA (Grozhik et al., 2019; Safra et al., 2017) and 2 – the mRNA coding for the PRUNE protein (Zhou et al., 2019). To examine the previously reported m1A sites on these RNAs, we used gene-specific RT-PCR to generate ∼500bp amplicons covering the regions of interest in both targets. This strategy enabled specific amplification of these less RNAs upstream of the NEXTERA library preparation that we used for all RNAs in this study. Using this method, we successfully obtained Mg/Mn mutational profiles for both RNAs in different cell lines (Figure 6). In the MALAT1 case, we observed a single nucleotide site with mutation rate roughly two orders of magnitude higher than other sites in both Huh7.5 and Hela cells (Figure 6A, B), which is on par with the range observed on 28S rRNAs used in the training set. The MRT-Modseq pipeline successfully identified this site as m1A-modified, confirming previous reports using different sequencing-based pipelines (Grozhik *et al*., 2019; Zhou *et al*., 2019). Similarly, the known m1A site in PRUNE1 mRNA exhibited a mutation rate significantly above those of other sites in both cell types (Figure 6C, D), albeit lower than that of the MALAT1 site. The relatively lower mutation rate in this case may reflect a lower modification stoichiometry, consistent with previous analyses showing that only up to ∼40% of PRUNE1 mRNAs are m1A-modified (Zhou *et al*., 2019). Importantly, these results demonstrate that MRT-Modseq can accurately flag modifications in RNAs that are far less abundant than rRNAs and even sub-stoichiometrically modified.

**Figure 6.**
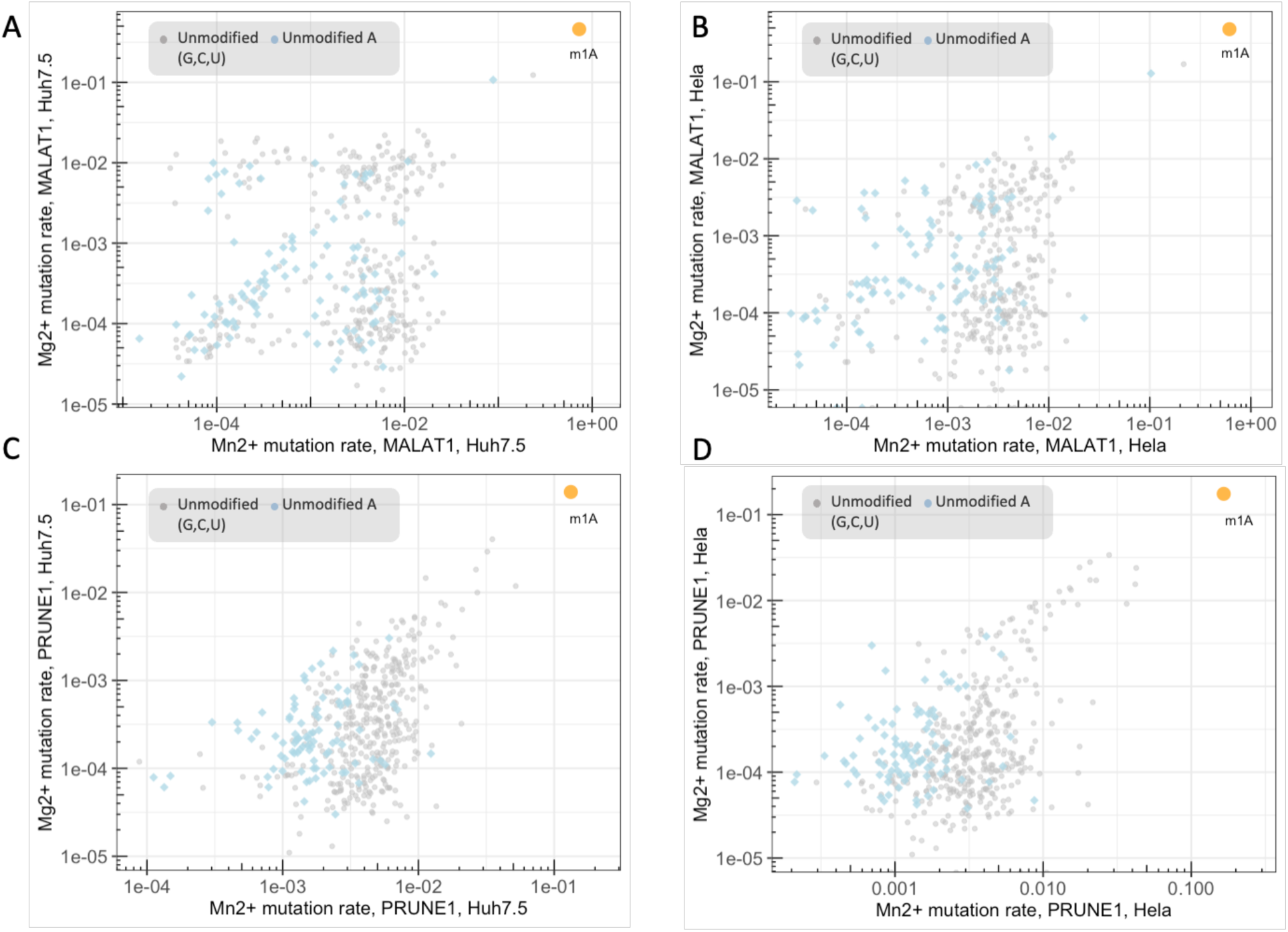
MRT-Modseq successfully captures known m1A sites in non-rRNA targets. (A) Log-scale plot of Mg (y-axis) and Mn (x-axis) mutation rates obtained with MRT for MALAT1 in Huh7.5 cells. Different symbols represent each group of nucleotides (unmodified A or unmodified G, C, U) as indicated in the legend. The known m1A site is represented by a yellow dot and is labelled below. (B) Log-scale plot of Mg (y-axis) and Mn (x-axis) mutation rates obtained with MRT for MALAT1 in Hela cells, labelled as in (A). (C) Log-scale plot of Mg (y-axis) and Mn (x-axis) mutation rates obtained with MRT for PRUNE1 in Huh7.5 cells, labelled as in (A). (D) Log-scale plot of Mg (y-axis) and Mn (x-axis) mutation rates obtained with MRT for PRUNE1 in Hela cells, labelled as in (A).

Taken together, our benchmarking results show that the MRT-ModSeq was highly efficient at detecting base modifications using experimental mutation rates and managed to correctly capture most sites of m7G, m1A, m3U, m1acp3Y and Nm modifications in the tested RNAs. Finally, our machine learning workflow showed robust performance in detecting sites of 2’-O-methylation after limited training on a prototype set of human rRNAs, which suggests the large potential and applicability of this approach as a platform for high-throughput screening of RNA modifications using MRT.

## DISCUSSION

The study of RNA modifications is critical to understanding RNA function in the cell, and yet it is fundamentally limited by our ability to map the identity and location of different deposited chemical groups in RNA transcripts at nucleotide resolution. Here, we present a new strategy to detect RNA modifications based on the reverse transcription signatures produced by MRT (MarathonRT). MRT-ModSeq does not require any chemical treatment of the RNA sample, a hallmark of several traditional RNA modification detection protocols. Instead, it is based on the differential rates of cDNA misincorporation of MRT in the presence of Mn^2+^ and Mg^2+^ (the “mutational signature”). We used the experimental signatures of RNAs with validated modification landscapes to train a supervised machine learning pipeline for detection and classification of RNA modifications. This method provides a rapid tool for identifying modification “hotspots” across a transcript and has the potential to significantly speed up discovery of modified RNAs in complex samples. We show that precision of the method is largely limited by the diversity of available RNA training sets, indicating that second-generation applications will provide even broader coverage of modification subtypes.

Traditionally, RT mismatch-based detection of RNA modifications has been achieved by computing mismatch rates from sequencing data and using the resulting signature as diagnostic for the presence of a modification (Li *et al*., 2017; Tan et al., 2021; Zhou *et al*., 2019). These methods usually do not involve classification, i.e., distinguishing between different modifications possible at the same nucleotide site. In general, high mutation rates observed in response to a particular type of modification cannot unambiguously serve as the sole diagnostic feature of that particular modification, since other (related) types of modification could in principle produce similar signals. In addition to detecting modified sites as positions of high mutation rates, the machine learning framework in MRT-ModSeq allows classification of different types of modifications based on the signatures present in the training data. This is possible because of the inherently multidimensional character of MRT mutational data (total and marginal mutation rates, insertions, deletions, etc), which is further expanded by the use of multiple enzymatic cofactors in our pipeline. Specifically, by employing an alternative divalent cation (Mn) for reverse transcription in addition to the canonical cofactor (Mg), we show that it is possible to use the mutational signatures to detect not only nucleobase modifications that directly affect Watson-Crick base pairing such as m1A, but also ribose 2’-OH methylations (Nm), which are otherwise silent under standard RT conditions (Motorin and Marchand, 2021) and therefore not amenable to this strategy of detection. Importantly, the unique ability of MRT to detect Nm sites in the presence of manganese effectively expands the conventional scope of detectable modifications by this strategy and enabled us to dissect the individual signatures of each class of Nm (Am, Gm, Cm, Um) sites.

Perhaps the most useful feature of MRT-ModSeq is that it can be carried out during the course of SHAPE-MaP studies that are aimed at secondary structure probing of natural RNAs in cells, lysates or from purified samples (Guo *et al*., 2020; Smola et al., 2015). By directly using an “untreated” RNA sample which serves as the negative control for SHAPE-MaP experiments and posterior structure prediction, one can easily implement the MRT-ModSeq cDNA workflow in parallel and identify endogenous modification sites at the same time as obtaining the secondary structure. In this way, MRT-ModSeq represents a simple, value-added way to expand the information obtained from conventional chemical probing experiments on natural RNA molecules.

We found that MRT signatures are remarkably variable relative to the type of RNA modification and their location in the templating nucleotide. Nucleobase modifications like m1A, m3U and m1ac3pY elicit the highest rates of misincorporation among all types analyzed, likely by directly interfering with canonical WC pairing and promoting mismatches at higher frequency relative to unmodified nucleotides. Modifications in other positions like the sugar ring (Nm) or the Hoogsteen edge (m7G) do not directly impede canonical WC pairing but they still appear to alter important interactions with the RT active site. The increased Mn mismatch rates were sufficient to enable detection, either by using a mutation rate cutoff (m7G) or via machine learning algorithms (Nm). Finally, RNA modifications like m5C, m6A and ψ were found to be mostly silent to MRT in both Mg and Mn conditions tested here, suggesting less pronounced effects on RT fidelity, even though they might lead to slight changes in cDNA misincorporation preferences as shown by our grouped ψ signature analysis (Fig 3). In these cases, alternative reaction conditions or RNA chemical treatment methods used in tandem with MRT-ModSeq (such as CMCT derivatization for ψ) may broaden the scope of positionally and chemically-assignable modifications within long transcripts.

In our analysis of the mismatch composition of modified nucleotides, we observed different nucleotide biases during cDNA misincorporation for different Nm groups, suggesting that the presence of a 2’-O-methyl group affects the conformational possibilities of each type of templating nucleotide and leads to specific selection of the preferred mismatch types that are accommodated by the RT active site. Here, it is helpful to look closer into the highly conserved substrate binding interface of group II intron reverse transcriptases (Lentzsch et al., 2021), as it includes a series of polar interactions with the RNA template, among which is a strong hydrogen bond between the 2’-OH group of the templating nucleotide and the peptide backbone (Stamos et al., 2017). One possibility is that the absence of this constraining interaction at 2’-O-methylated sites and the extra bulkiness of the methyl group are likely to disturb that interface and enhance sampling of alternative conformations, thereby favoring the occurrence of cDNA misincorporations. These effects are further enhanced by the use of manganese as a cofactor, which is routinely employed in RNA structure probing experiments to map 2’-OH adduct positions using RT mutation rates (Siegfried et al., 2014; Smola *et al*., 2015). Structural studies of MRT in complex with modified RNA substrates will be important to elucidate the interplay between the observed signatures for each type of modification and the enzymatic determinants of base pairing fidelity.

Interestingly, we noticed a general absence of significant sequence context effects on the signatures produced by MRT at Nm and ψ nucleotides, as revealed by our analysis of marginal mutation rates as a function of the penultimate nucleotide (Fig 4). This verification obviated the inclusion of any explicit sequence-based metrics in our machine learning algorithms as classification features. Our observations are consistent with studies using other reverse transcriptases that detected no evident sequence biases in cDNA synthesis of RNA templates containing pseudouridines and other base modifications (Potapov et al., 2018). Since the RT signatures of modifications like m1A have been shown to be context-dependent in previous studies (Hauenschild et al., 2015), it remains to be assessed whether this is also the case with other types of base modification (e.g., m3U, m7G, etc), which will be important in future implementations of MRT-ModSeq specifically designed to detect them.

When compared to other methods that also employed machine learning for Nm detection, MRT-ModSeq achieves >99% accuracy across nucleotide types, which is equivalent or superior to most available ML-based methods in the literature (Chen et al., 2016; Mostavi et al., 2018; Qiu et al., 2017; Tahir et al., 2019; Yang et al., 2018). In addition, the detection accuracy of MRT-ModSeq is largely comparable to that of a dedicated experimental pipeline such as 2OMe-seq (Incarnato et al., 2017), which further illustrates the power of using MRT signatures to detect specific types of RNA modifications.

MRT-ModSeq benchmarking results provide important clues on the functioning and performance of machine learning algorithms trained on a limited prototype dataset of human 18S and 28S rRNAs. Out of the 4 ML algorithmic prongs used for Nm detection, Am and Gm (average MCC = 0.87, in both cases, Fig S10) showed the best performance, when compared to Um and Cm (average MCC = 0.79 and 0.78, respectively, Fig S10). Interestingly, Am and Gm nucleotides produced the strongest signatures among Nm groups, as revealed by our group analysis of marginal mutation rates (Fig 3), suggesting a correlation between the magnitude of mutational signatures used for training and the performance of ML detection. Given these observations, we predict that highly mutagenic modification types like the ones we detected using mutation rate cutoffs (m1A, m7G, m3U, etc) are likely to result in high performance of ML detection once appropriate training sets are employed that sample a large diversity of these sites. In addition to rRNAs, we also tested MRT-ModSeq on selected RNAs with experimentally-validated modifications and showed that the pipeline readily detects known m1A sites on both MALAT1 lncRNA and PRUNE1 mRNA, despite their reduced abundance and modification stoichiometry. The targeted approach we used in this case (i.e., RT-PCR amplification on polyA-enriched RNA) can be conveniently used to detect specific modified sites in other mRNAs and lncRNAs, and will hopefully be a valuable tool to confirm the presence of individual modifications in studies testing their functional roles.

On the other side of the spectrum, modifications such as pseudouridines (ψ), which displayed only minor signatures relative to unmodified uridines, showed inferior performance on MRT-ModSeq benchmarking tests (average MCC ∼0.28). The reduced detection power in this case can be attributed to the minor perturbation introduced by pseudouridine during MRT reverse transcription, since canonical WC base pairing is not affected by pseudouridylation. Future work exploring reaction conditions will be important to identify factors that could sensitize MRT towards ψ sites and then to evaluate the impact of improved mutation segregation on ML classification performance for both pseudouridines and also other modifications that were mostly mismatch-silent under ModSeq conditions, like m6A and m5C (Fig 1). However, ψ detection may represent a class of modifications that will ultimately require a complementary sequencing approach (Carlile et al., 2014; Khoddami et al., 2019; Schwartz et al., 2014) that is used in tandem with ML ModSeq approaches for classification, and these can be combined in future studies to increase detection performance.

### Limitations of the study

The sensitivity of MRT-ModSeq in its current form is limited by the training set used for assigning the identity of chemical modifications. The modification landscape within the ribosomal RNAs used in this study constitutes only a small subset of the many important modifications within cellular RNAs. For example, in the system we used for Nm classification, it remains possible for other types of RNA modification to display signatures that overlap with those of Nm sites. This is possible because other modifications not included in the original training data could have a mutational fingerprint that overlaps with the ones used as the training set. The findings reported here underscore the importance of expanded training sets based on additional synthetic and natural modified RNAs, along with orthogonal methods (e.g., mass spectrometry (Lauman and Garcia, 2020), enzymatic (Mikutis et al., 2020), IP-based (Dominissini et al., 2016)).

Another limitation of our pipeline is that, in its current state, it cannot retrieve quantitative information on modification levels or stoichiometry. However, this is an interesting possibility for future methodological development, which will likely involve fine-tuning RT reaction conditions and then assessing the effects on the sensitivity of detection, in addition to the generation of new machine learning parameters to enable quantification. Although we focused the initial development of MRT-ModSeq on the screening of RNA modifications in specific targets, an attractive possibility is its application to transcriptome-wide profiling. With the due optimization of sample preparation and computational parameters, we anticipate that future MRT-ModSeq versions will enable the unbiased discovery of modified sites across several transcripts in a single sequencing experiment.

### Significance

In the present work, we introduce MRT-ModSeq, a general platform trained on curated RT fingerprints to detect multiple RNA modifications in a single pipeline. Since MRT-ModSeq does not require dedicated RNA sample pre-treatment or modification-specific steps, it is particularly useful in cases where there is little or no information available on the positions or identities of chemical modifications across the RNA(s) of interest (e.g., novel targets, new biological samples, different cellular states, etc.), and it can be used in parallel with conventional secondary structure determination workflows. The method is general in that the set of RNAs used for training can be as large and diverse as practically feasible, which includes the possibility of both natural and synthetically modified RNA libraries. This strategy is applicable not only to abundant RNAs such as the ones we utilized for training and testing, but also to lower abundance transcripts through the use of gene-specific RT-PCR amplification, as shown in our analysis of PRUNE1 mRNA and MALAT1 lncRNA. By employing a suitable library of training RNAs, MRT-ModSeq can be purposefully adapted to the detection of new types of RNA modifications that extend beyond the ones we describe in this work. Finally, we anticipate that the superior processivity of MRT will enable coupling of the ModSeq methodology with long-read sequencing in order to profile chemical modifications at single molecule resolution.

## DATA AND CODE AVAILABILITY

All scripts used for data parsing and filtering were written in the R computing language. The complete MRT-ModSeq computational package including documentation is publicly available for use on the Pyle Lab GitHub repository (https://github.com/pylelab/MRTModSeq). The data used in this publication have been deposited in NCBI’s Gene Expression Omnibus (Edgar et al., 2002) and are accessible through GEO Series accession number GSE202160 (https://www.ncbi.nlm.nih.gov/geo/query/acc.cgi?acc=GSE202160).

## ACKNOWLEDGEMENTS

This work was supported by the Howard Hughes Medical Institute (A.M.P.); the National Human Genome Research Institute (R01 HG011868-01 to A.M.P.) and a Yale College Dean’s Research Fellowship (to G.M.). Funding for open access charge was provided by the Howard Hughes Medical Institute. The authors thank members of the Pyle lab at Yale University, especially Dr Li-Tao Guo and Dr Chengxin Zhang, for insightful discussion of the results and methodology reported in this work. The authors also thank Dorthy Fang from Dr Sigrid Nachtergaele lab for useful experimental suggestions.

## AUTHOR CONTRIBUTIONS

R.T., G.M. and A.M.P. conceived the project. R.T. and H.W. designed and performed the experiments. G.M. developed the MRT-ModSeq computational pipeline and analyzed the benchmarking dataset. H.W. analyzed data on MALAT1 and PRUNE1. All authors contributed to writing and editing of the manuscript.

## DECLARATION OF INTERESTS

The authors declare no competing interests.

## INCLUSION AND DIVERSITY STATEMENT

We support inclusive, diverse, and equitable conduct of research.

## METHODS

### Cell culture

Huh7.5, HeLa (ATCC CCL-2) and HEK293 (ATCC CRL-1573) cell lines were maintained in Dulbecco’s Modified Eagle Medium (Invitrogen 11965092) supplemented with 10% fetal bovine serum (Invitrogen 16140071). The Huh7.5 cell line was a gift from Dr Brett Lindenbach (Yale University). For total RNA extraction and library preparation, cells were seeded in 10-cm plates and grown until ∼80% confluent.

*Escherichia coli* (NEB 5-alpha, NEB C2987H) were grown in 5 ml LB broth at 37°C and shaking at 180 rpm until the culture reached OD_600_ 0.7. Cells were pelleted by centrifugation at 6,000 x g for 15 min at 4°C and immediately used for total RNA extraction.

*Saccharomyces cerevisiae* wild-type strain (NB40-36a) was kindly provided by T. Fox (Perez-Martinez et al., 2003). *S. cerevisiae* were grown from single colony in 5 ml of YPD medium overnight (30°C, 330 r.p.m.). The following day, a 1 ml volume of cultured cells was pelleted by centrifugation at 3000 r.p.m. for 1 min and immediately used for total RNA extraction.

### Total RNA extraction

For RNA extraction from mammalian cell lines, cells at ∼80% confluency (10-cm plates) were first washed with DPBS (Genesee 25-508), then trypsinized and pelleted by centrifugation at 200 x g for 5 min at 4°C. Cell pellets were resuspended in 1 ml TRIzol reagent (Invitrogen 15596018) and incubated at room temperature for 5 min. RNA was isolated from the TRIzol lysate according to the manufacturer’s guidelines. Briefly, the lysate was treated with chloroform, followed by centrifugation at 12,500 x g for 15 min. The aqueous phase was then purified on a Zymo RNA clean and concentrator column (Zymo R1013). RNA samples were treated with RQ1 RNase-Free DNase (Promega M6101) and cleaned up using a Zymo RNA clean and concentrator kit.

For RNA extraction from bacterial cells, the *E. coli* cell pellet was resuspended in 200 ul of Max Bacterial Enhancement Reagent (Invitrogen 16122012) and incubated at 95°C for 4 min. The sample was then dissolved in 1 ml TRIzol and incubated for 5 min at room temperature. Total RNA was then extracted from the TRIzol lysate, DNAse-treated and purified using the same procedure as for mammalian cells.

For RNA extraction from yeast cells, the yeast cell pellet was washed twice with water and lysed in 700 ul TRIzol + 300 ul acid-washed glass beads (100 um) while vortexing at top speed for 1 minute. Total RNA was isolated from the TRIzol lysate, DNAse-treated and purified as described for mammalian cells.

### MRT reverse transcription (cDNA synthesis)

MRT (MarathonRT) used in all experiments was expressed and purified according to the standard, optimized protocol reported in Guo et al. 2020 (Guo *et al*., 2020). Total RNA from each biological sample was reverse-transcribed using MRT in two separate reactions in the presence of either magnesium or manganese as cofactors. Total RNA (2 µg) was first annealed with 200 ng random nonamer primers (NEB S1254S) by incubation at 70°C for 5 min, then at 4°C for 2 min. Reverse transcription was then performed in optimized MRT buffer (Guo et al. 2020) containing 50 mM Tris–HCl (pH 7.5), 200 mM KCl, 5 mM DTT, 20% glycerol, 0.5 mM dNTP mix, in the presence of either cofactor (1 mM MgCl_2_ or 2 mM MnCl_2_) and 2 ul of MRT (40 U), to a final reaction volume of 20 ul. The reaction was incubated at 42°C for one hour, followed by enzyme inactivation at 70°C for 15 min. The cDNA sample obtained in this manner was immediately used for subsequent Nextera XT library preparation.

### Sequencing library construction

Nextera XT libraries were built from cDNA obtained in each MRT reaction (Mg or Mn). First, second strand synthesis was performed with the NEBNext Ultra II Non-Directional Second Strand Synthesis Module (NEB E6111S) following the manufacturer’s instructions. The dsDNA was then purified with the Monarch PCR and DNA cleanup kit (T1030S) and quantified on an Invitrogen Qubit fluorometer. Paired-end sequencing libraries were generated from each dsDNA sample using the Nextera XT DNA Library Preparation Kit (Illumina FC-131-1024) exactly as recommended by the manufacturer. Final libraries were purified with AMPure XP beads (Beckman Coulter A63880) using a 1:1 beads/sample ratio, quantified on an Invitrogen Qubit fluorometer and analyzed on an Agilent 2100 Bioanalyzer. Libraries were sequenced on an Illumina NextSeq 500 instrument.

### MRT-ModSeq tests on MALAT1 lncRNA and PRUNE1 mRNA

#### PolyA enrichment

RNA extracted as described above from HeLa and Huh7.5 cell lines was subjected to polyA fraction enrichment prior to MRT-ModSeq library preparation. Dynabeads MyOne Streptavidin C1 (Invitrogen 65001) were first incubated with Biotinylated Oligo (dT) probes (20mer, ThermoFisher) at 42°C for 30 min in hybridization buffer (10 mM Tris–HCl pH 7.5, 1 mM EDTA, 500 mM LiCl, 4M Urea, 2.5 mM TCEP). Beads were washed twice on a magnet with hybridization buffer and then incubated with total cellular RNA at 42°C for 2 hours in hybridization buffer with intermittent mixing at 1000 rpm (thermomixer). For 40ug of total cellular RNA input, 200 pmol Biotinylated probe and 40 ul of beads were used. After hybridization, beads were washed twice with hybridization buffer and RNA was eluted in water by heating the beads at 95°C for 1 min. Enriched RNA was then cleaned up using SILANE beads (Invitrogen 37002D) and used for MRT reverse transcription.

#### Reverse transcription and amplicon PCR

MRT reverse transcription was performed as described above but using 1 pmol of gene-specific primer per reaction. Following reverse transcription, cDNA was cleaned up with AMPure XP beads (Beckman A63881) and amplified with Q5 Hot Start Polymerase (NEB M0491S). RT-PCR primers used are listed below.

MALAT1 RT: CTTCTCAAAACACCAGCTG

MALAT1 PCR_F: CAGGTGAACATAACAGACTTGG

MALAT1 PCR_R: CTCAAAACACCAGCTGCAGG

PRUNE1 RT: TTTTGGATAAGATATGATG

PRUNE1 PCR_F: CAGTTCCTCCCGGGGTCG

PRUNE1 PCR_R: TGGATAAGATATGATGGTCGAC

### Sequencing data alignment and computation of input features for machine learning

Sequencing data were aligned to reference RNA sequences using ShapeMapper (Smola *et al*., 2015). ShapeMapper uses two sets of paired-end sequencing data (untreated and treated). For use as inputs to ShapeMapper, we designated Mg libraries (reverse transcription with magnesium) the ‘untreated’ sample and Mn libraries (reverse transcription with manganese) the ‘treated’ sample. Fastq files corresponding to both samples were aligned to each individual RNA sequence (fasta format) with the Bowtie2 implementation in the ShapeMapper script and sequencing stats (depth, mutation counts, mutation rates) were outputted for each nucleotide position. The ‘--output-counted-mutations’ flag on ShapeMapper was used to output mutational composition rates (insertions, deletions, individual mismatches, etc) that were used as machine learning input features, i.e., the measurable properties that describe any given nucleotide site on the RNA transcript. Reproducibility was assessed by correlating mutation rates across two independent biological experiments in all cases.

### Numerical criteria for determining the presence of nucleobase modifications

Consistent with the signatures detected for human 18S and 28S rRNAs in the training set, uridine sites with a total Mn^2+^ mutation rate above 0.5 were classified as a “m3U or m1acp3Y”. Adenosine sites with a total Mn^2+^ mutation rate above 0.5 were classified as a “m1A”. Guanosine sites with a total Mn^2+^ mutation rate above 0.05 and total Mg^2+^ mutation rate above 0.0015 were classified as “m7G”. Cutoff values were chosen to minimize the overlap with unmodified sites, so that nucleotides initially identified by these filters are strong candidates for posterior validation. Only nucleotides that did not meet the above cutoffs were further employed in the downstream machine learning workflow.

### Data filtering and pre-processing

For each pair of Mg and Mn libraries, a series of mutational composition rates, along with overall mutation rates, were computed with ShapeMapper, in addition to nucleotide identity, nucleotide position and effective read depth. In total, 24 mutational composition features (predictors) were computed for each RNA, comprising all possible changes in nucleotide identity relative to the reference sequence for each site: total mutation rates, simple insertion/deletion rates, and complex insertion/deletion rates in both experimental conditions (Mg and Mn RT reactions). Mutational composition features were normalized against the total mutation rate for a given site on the transcript (all composition features sum to 1). The true modification status was appended to the datasets for each nucleotide site. Training datasets (18S rRNA Mg & Mn, 28S rRNA Mg & Mn) were combined to build the final training set for machine learning models. The combined dataset was first filtered to ensure read depth >5000 for each nucleotide site across the transcripts and only positions meeting this criterion were used for downstream analysis, which still accounted for >90% of all nucleotides in the dataset. The dataset was then split into four subsets by nucleotide type. Prior to training, positions containing rare modifications and/or nucleobase modifications (m1acp3Y, m6A, ac4C, m1A, pUm, m3U, m6Am, m7G and m5C) were filtered out of the dataset. We sorted and streamlined the features by conducting zero/low-variance tests to ensure robust performance of forest-based algorithms: from the original 24 parameters outputted by ShapeMapper, a zero-variance test (Kuhn, 2008) was used to pare down the features to the final 9 features used in ML models; namely, multi-nucleotide insertion/deletion and complex insertion/deletion mutation rates in the presence of Mg^2+^ or Mn^2+^ were removed, along with the extraneous mutational composition rates with initial identities not matching the canonical nucleotide identity. The final features list included: Mn mutation rate (‘Mn_rate’), Mg mutation rate (‘Mg_rate’), Mn single deletion rate (‘Mn_X-‘), and six mutational composition rates specific to the nucleotide type (e.g., for a site with canonical nucleotide A, ‘Mn_AT’, ‘Mn_AC’, ‘Mn_AG’, ‘Mg_AT’, ‘Mg_AC’, ‘Mg_AG’).

### Supervised machine learning workflow

To classify Nm and ψ sites on a transcript using MRT-ModSeq data, we implemented a supervised learning pipeline. All components of the algorithm training and testing – feature selection, resampling, training and cross-validation, benchmarking, and testing – were implemented in the Java-based Waikato Environment for Knowledge Analysis (WEKA) framework (Smith and Frank, 2016). All training and testing based on human ribosomal RNAs was conducted using reference datasets obtained from Huh7.5 total cellular RNA. In all training and benchmarking routines, only mapped sites with read depth >5000 were used. Consistently, the number of modified and unmodified sites reported in benchmarking results (Tables S1-S7) add up to the total number of mapped sites in MRT-ModSeq data with sufficient sequencing coverage.

All tests for class balancing (undersampling, oversampling, hybrid) were conducted through WEKA’s synthetic minority oversampling implementation (Chawla *et al*., 2002). For the final model training, the original datasets were used in all cases. We benchmarked Naïve Bayes, k-Nearest Neighbor, Support Vector Machine, Random Forest, and Neural Network algorithms, along with bagged and boosted versions of each algorithm (15 algorithms in total) using WEKA’s Experimenter Workbench. Benchmarking was conducted on each training set in the resampling loop using an inner 10-fold cross validation loop. This process was repeated for each nucleotide type. Kappa statistic, Matthews Correlation Coefficient (MCC), F-measure, and Accuracy (ACC) were used to score performance in each case. Importantly, nucleobase-modified sites (subject to classification via the high-pass filter) were not included within the training routine. The algorithms for A, C, and G are binary classification algorithms (between ‘unmodified’ and ‘Nm’), while the algorithm for U is a ternary classification algorithm (between ‘unmodified’, ‘Nm’, and ‘pseudoU’). Boosted tree-based methods (AdaBoost-RT) were used as final training algorithms for adenosines and cytidines, while bagged tree-based methods (Bagging-J48) were used as final training algorithms for guanosines and uridines. The unaltered output of the 10-fold cross validation training was denoted the ‘permissive algorithm’. To build the ‘stringent algorithm,’ we implemented a cost matrix that imparted a 10x cost to misclassification of unmodified sites as modified (false positives). This cost matrix was built into the 10-fold cross validation evaluation step in WEKA.

### Performance metric definitions

TP = number of true positives; TN = number of true negatives; FP = number of false positives; FN = number of false negatives

- Matthews Correlation Coefficient (Chicco and Jurman, 2020), MCC = 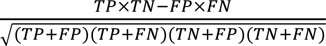
- Cohen’s Kappa Statistic (McHugh, 2012), 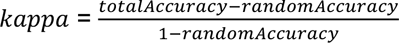, where 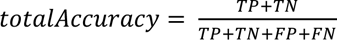, and 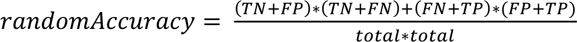
- F-measure (Chicco and Jurman, 2020), 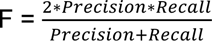, where 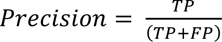 and 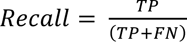
- Accuracy (Chicco and Jurman, 2020), 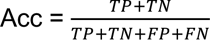

### Reference RNA sequences and ground-truth modification sites

Reference sequences for human 18S rRNA (1869 bp, NCBI ID: 100008588), human 28S rRNA (5070 bp, NCBI ID: 100008589) and human 5.8S rRNA (157 bp, NCBI ID: 100008587) were downloaded from the GenBank database (Benson et al., 2013). Ground-truth modification sites for the human ribosomal RNAs were extracted from Taoka et al. (Taoka *et al*., 2018). Sequences for *E. coli* 16S rRNA (1541 bp, GeneBank ID: J01859.1) and 23S rRNA (2904 bp, GeneBank ID: V00331.1) were downloaded from the GenBank database. Ground-truth modification sites for the *E. coli* ribosomal RNAs were obtained from the MODOMICS database (Boccaletto et al., 2022) and confirmed with a recent cryo-EM based study (Stojkovic et al., 2020). Reference sequences for *Saccharomyces cerevisiae* 18S rRNA (1800 bp, RNAcentral ID: URS00005F2C2D_4932) and 25S rRNA (3396 bp, RNAcentral ID: URS000061F377_4932) were downloaded from the RNAcentral database (Consortium, 2021). Ground-truth modification sites for the *Saccharomyces cerevisiae* ribosomal RNAs were obtained from the MODOMICS database and the assignments were further verified against a recent study cataloging the yeast rRNA modification landscape (Yang *et al*., 2016). Reference sequences for human MALAT1 (NR_002819.4) and human PRUNE1 (NM_021222.3) were obtained from GenBank.

## SUPPLEMENTAL INFORMATION

MRT-ModSeq_SUPPLEMENTARY_DATA.pdf

Figures S1-S10. Tables S1-S11.

## Notes

### Competing Interest Statement

The authors have declared no competing interest.

