## Supplementary material for "MRT-ModSeq – Rapid detection of RNA modifications with MarathonRT": Figures S1-S10 Tables S1-S11

**SUPPLEMENTARY DATA for *MRT-ModSeq – Rapid detection of RNA modifications with MarathonRT***

**This file contains:**

Supplementary Figures

| | |
|---|---|
| Fig. S1 | Related to Fig. 1 |
| Fig. S2 | Related to Fig. 1 |
| Fig. S3 | Related to Fig. 1 |
| Fig. S4 | Related to Fig. 1 |
| Fig. S5 | Related to Fig. 1 |
| Fig. S6 | Related to Fig. 1 |
| Fig. S7 | Related to Fig. 4 |
| Fig. S8 | Related to Fig. 4 |
| Fig. S9 | Related to Fig. 4 |
| Fig. S10 | Related to Fig. 5 |

Supplementary Tables

| | |
|---|---|
| Tables S1-S11 | MRT-ModSeq benchmarking results |

### SUPPLEMENTARY FIGURES

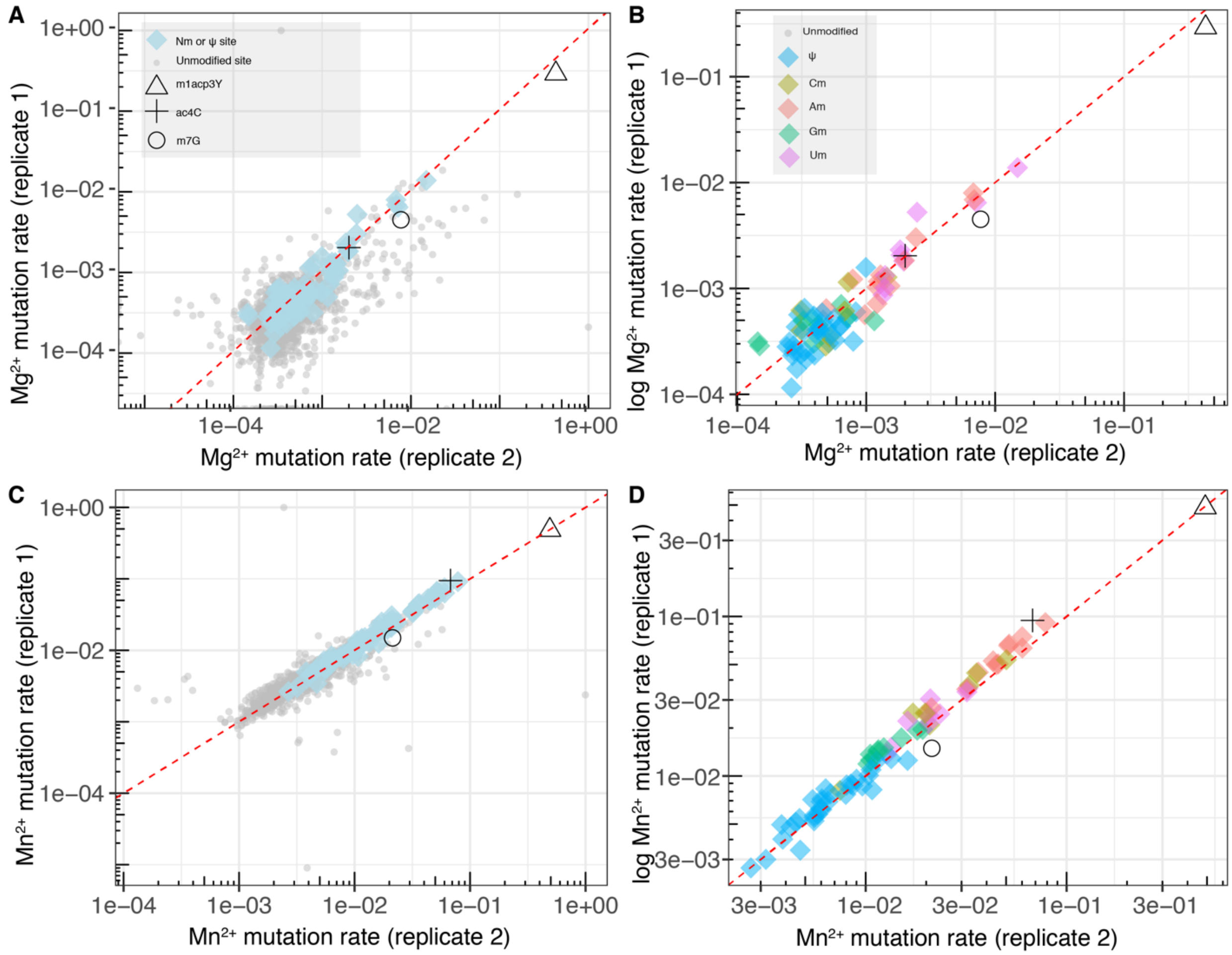


**Figure S1.** MRT mutation rate reproducibility across experimental replicates for the human 18S rRNA, sequenced from Huh7.5 cells. (A) Log-scale plots of Mg mutation rates for two independent experiments, for all nucleotide sites, with specific modification status shown in the legend on the top left-hand corner. (B) Similar plot as in (A), but only showing Nm and ψ positions. (C) Similar plot as in (A), but showing Mn mutation rate values. (D) Similar plot as in (B), but showing Mn mutation rates.

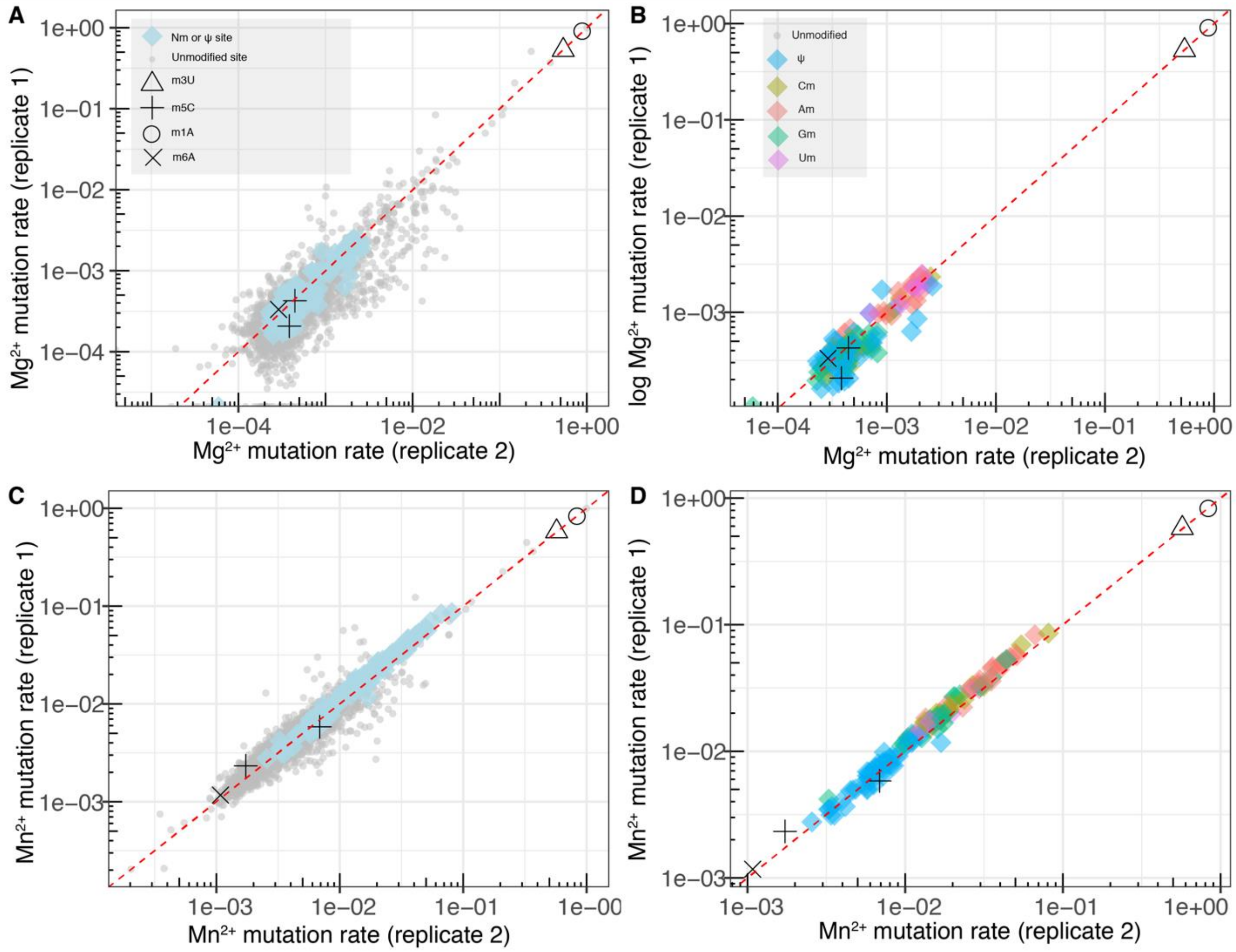


**Figure S2.** MRT mutation rate reproducibility across experimental replicates for the human 28S rRNA, sequenced from Huh7.5 cells. (A) Log-scale plots of Mg mutation rates for two independent experiments, for all nucleotide sites, with specific modification status shown in the legend on the top left-hand corner. (B) Similar plot as in (A), but only showing Nm and ψ positions. (C) Similar plot as in (A), but showing Mn mutation rate values. (D) Similar plot as in (B), but showing Mn mutation rates.

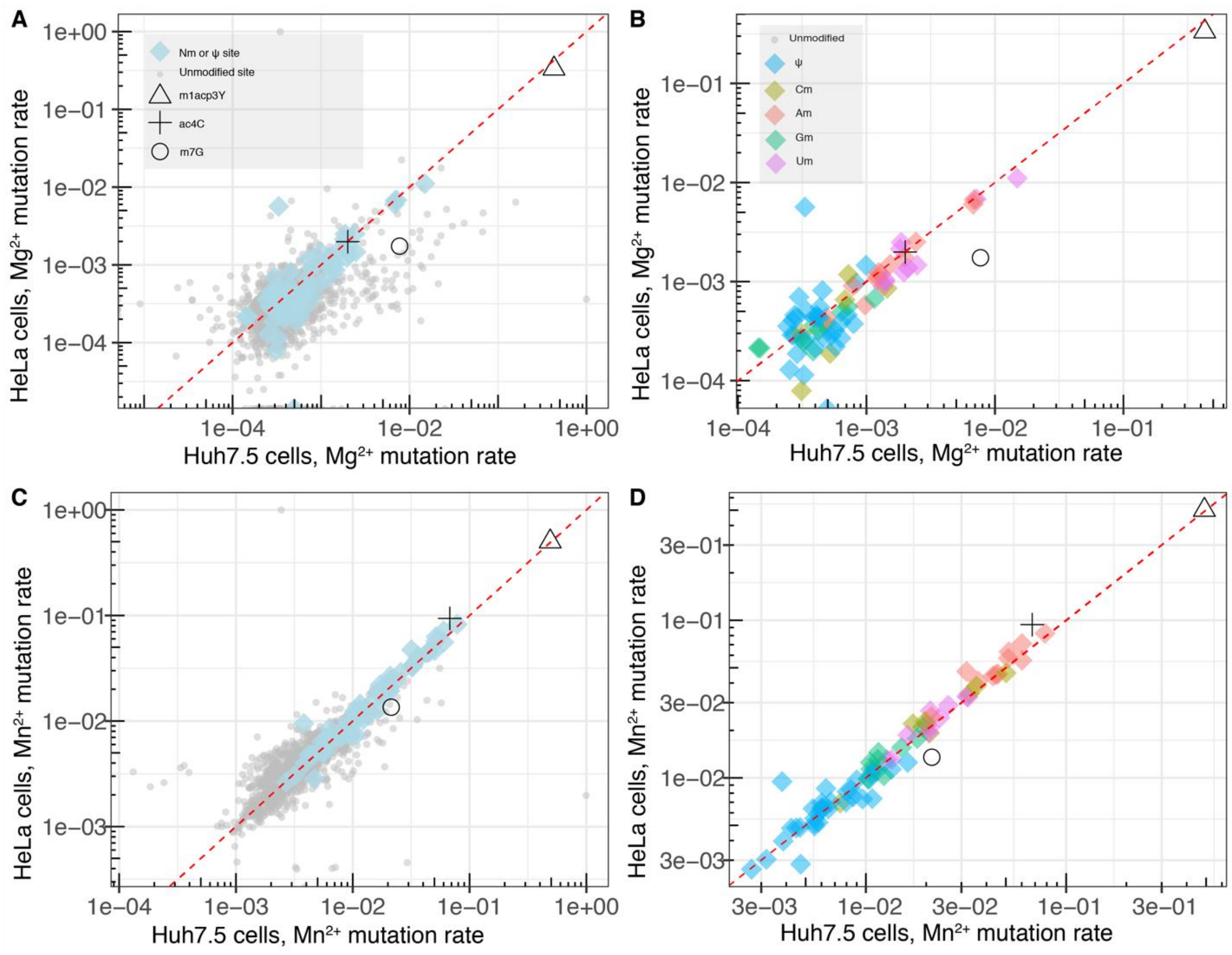


**Figure S3.** MRT mutation rate consistency between Huh7.5 and HeLa cell lines for human 18S rRNA. (A) Log-scale plots of Mg mutation rates calculated from HeLa (y-axis) and Huh7.5 (x-axis) datasets, for all nucleotide sites, with specific modification status shown in the legend on the top left-hand corner. (B) Similar plot as in (A), but only showing Nm and ψ positions. (C) Similar plot as in (A), but showing Mn mutation rate values. (D) Similar plot as in (B), but showing Mn mutation rates.

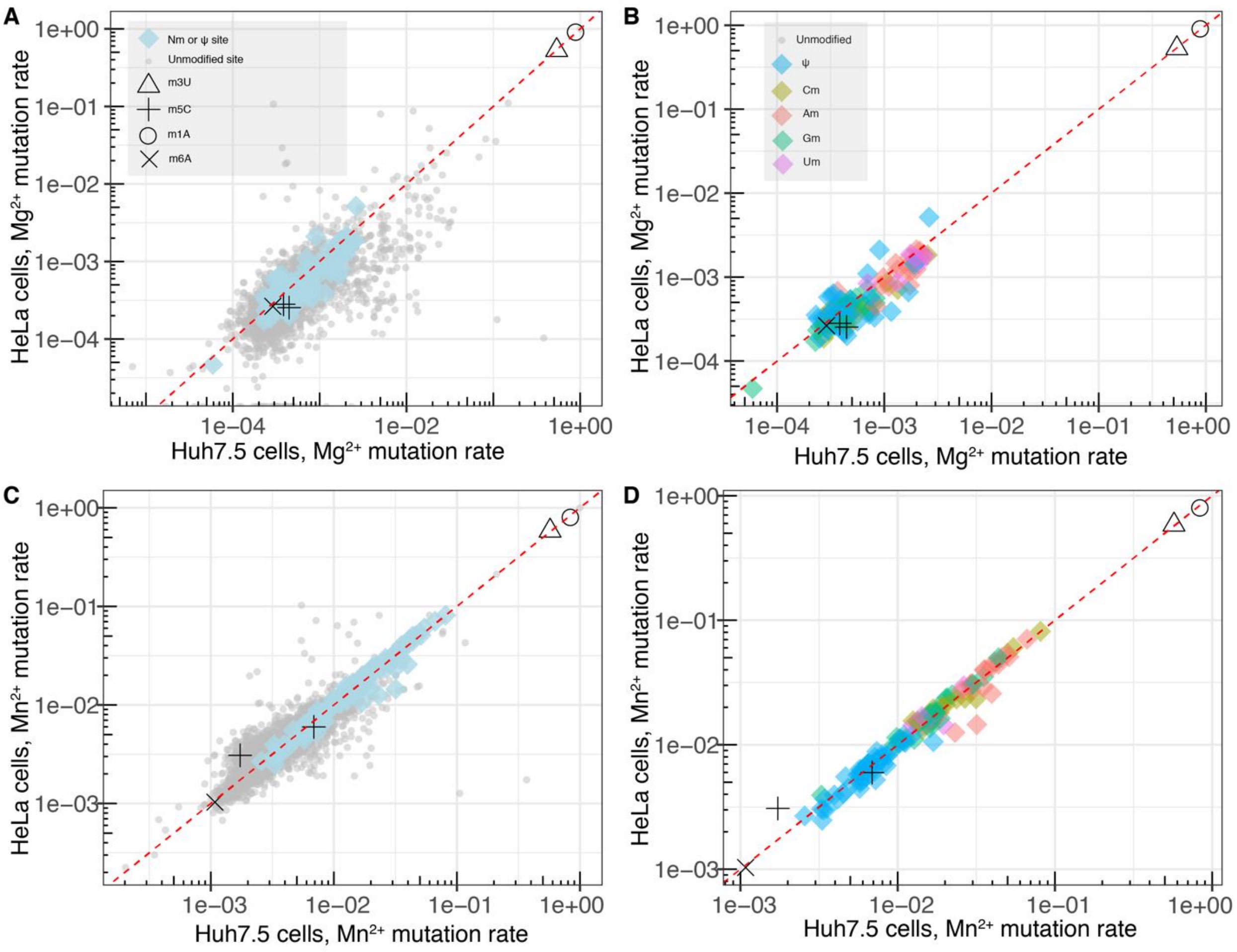


**Figure S4.** MRT mutation rate consistency between Huh7.5 and HeLa cell lines for human 28S rRNA. (A) Log-scale plots of Mg mutation rates calculated from HeLa (y-axis) and Huh7.5 (x-axis) datasets, for all nucleotide sites, with specific modification status shown in the legend on the top left-hand corner. (B) Similar plot as in (A), but only showing Nm and ψ positions. (C) Similar plot as in (A), but showing Mn mutation rate values. (D) Similar plot as in (B), but showing Mn mutation rates.

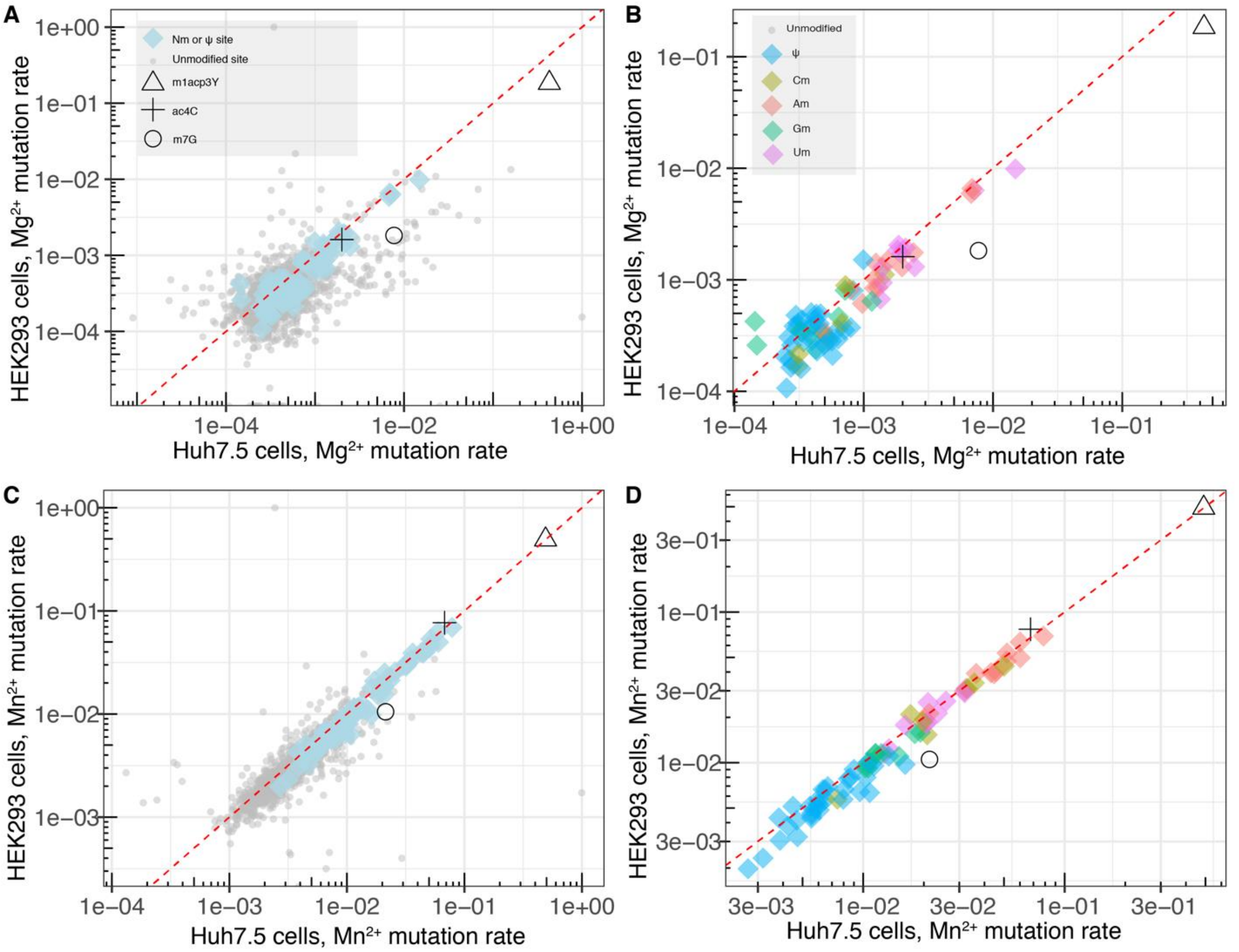


**Figure S5.** MRT mutation rate consistency between Huh7.5 and HEK293 cell lines for human 18S rRNA. (A) Log-scale plots of Mg mutation rates calculated from HEK293 (y-axis) and Huh7.5 (x-axis) datasets, for all nucleotide sites, with specific modification status shown in the legend on the top left-hand corner. (B) Similar plot as in (A), but only showing Nm and ψ positions. (C) Similar plot as in (A), but showing Mn mutation rate values. (D) Similar plot as in (B), but showing Mn mutation rates.

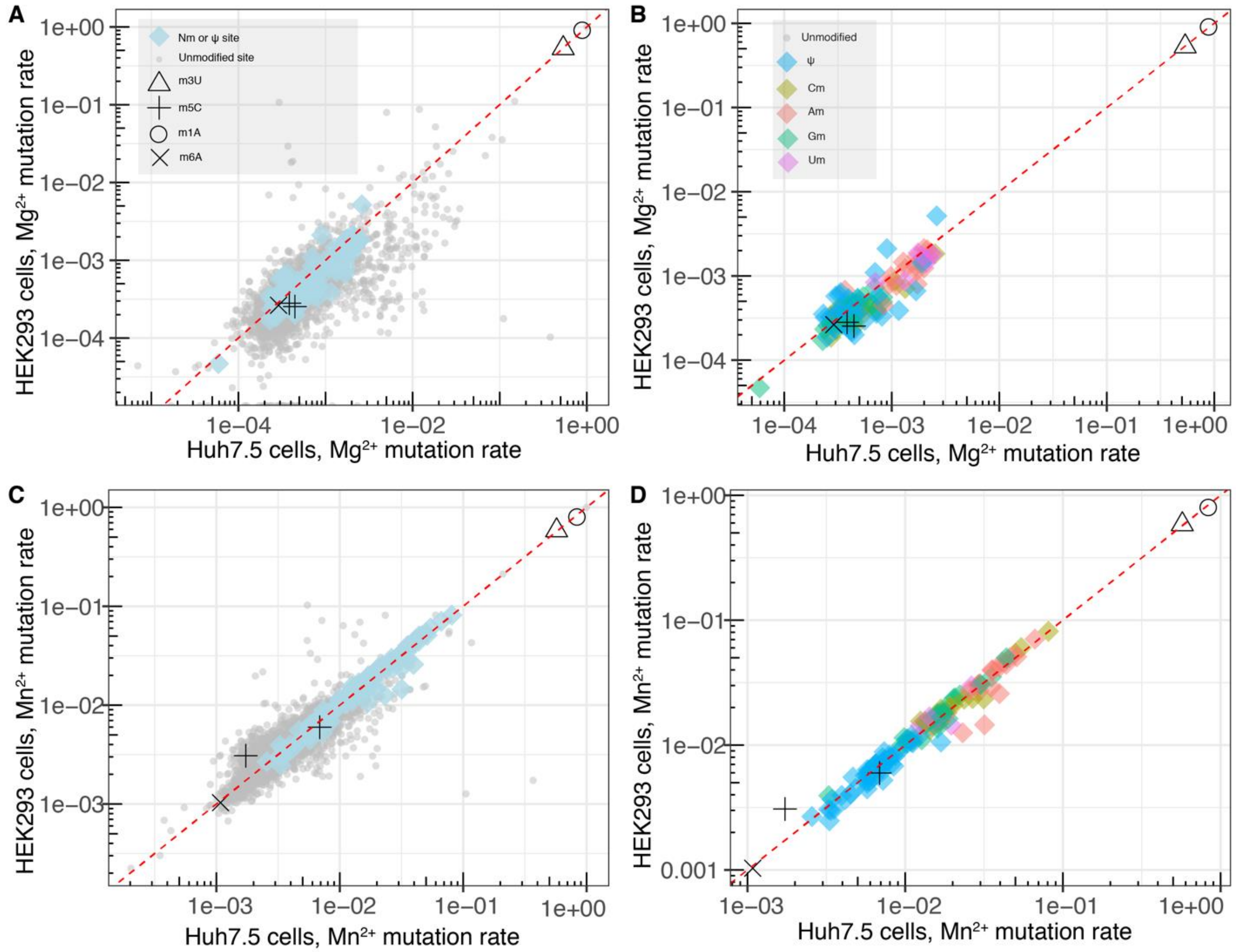


**Figure S6.** MRT mutation rate consistency between Huh7.5 and HEK293 cell lines for human 28S rRNA. (A) Log-scale plots of Mg mutation rates calculated from HEK293 (y-axis) and Huh7.5 (x-axis) datasets, for all nucleotide sites, with specific modification status shown in the legend on the top left-hand corner. (B) Similar plot as in (A), but only showing Nm and ψ positions. (C) Similar plot as in (A), but showing Mn mutation rate values. (D) Similar plot as in (B), but showing Mn mutation rates.

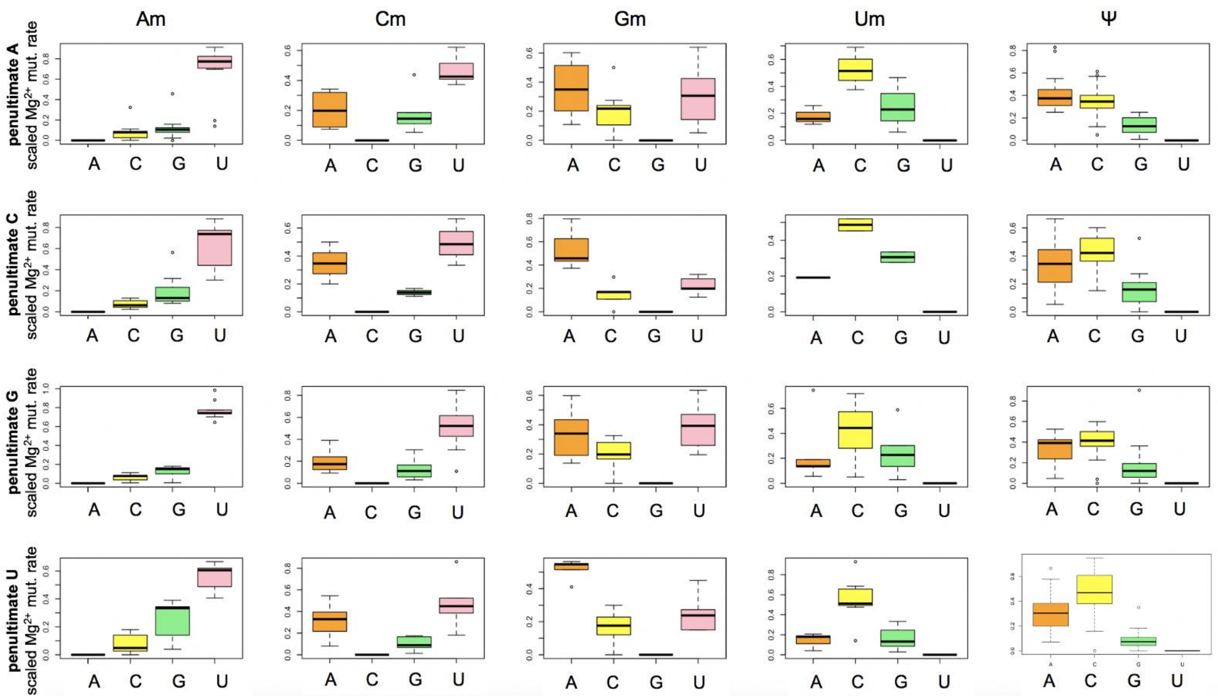


**Figure S7.** Sequence context analysis of magnesium (Mg) mutational signatures of Nm and ψ sites in the combined 18S/28S rRNA dataset. The results are sorted by modified nucleotide type (columns) and by the penultimate nucleotide in the RNA sequence, i.e., the nucleotide immediately 3' to the position being examined. Each individual plot shows the distribution of marginal (scaled) mutation rates calculated from the Mg sample as a function of the different groups of mutated nucleotides (e.g., the top graph in the Am column shows marginal mutation rates from A to each one of the 4 nucleotides: A to A (no mutation), A to C, A to G, A to U).

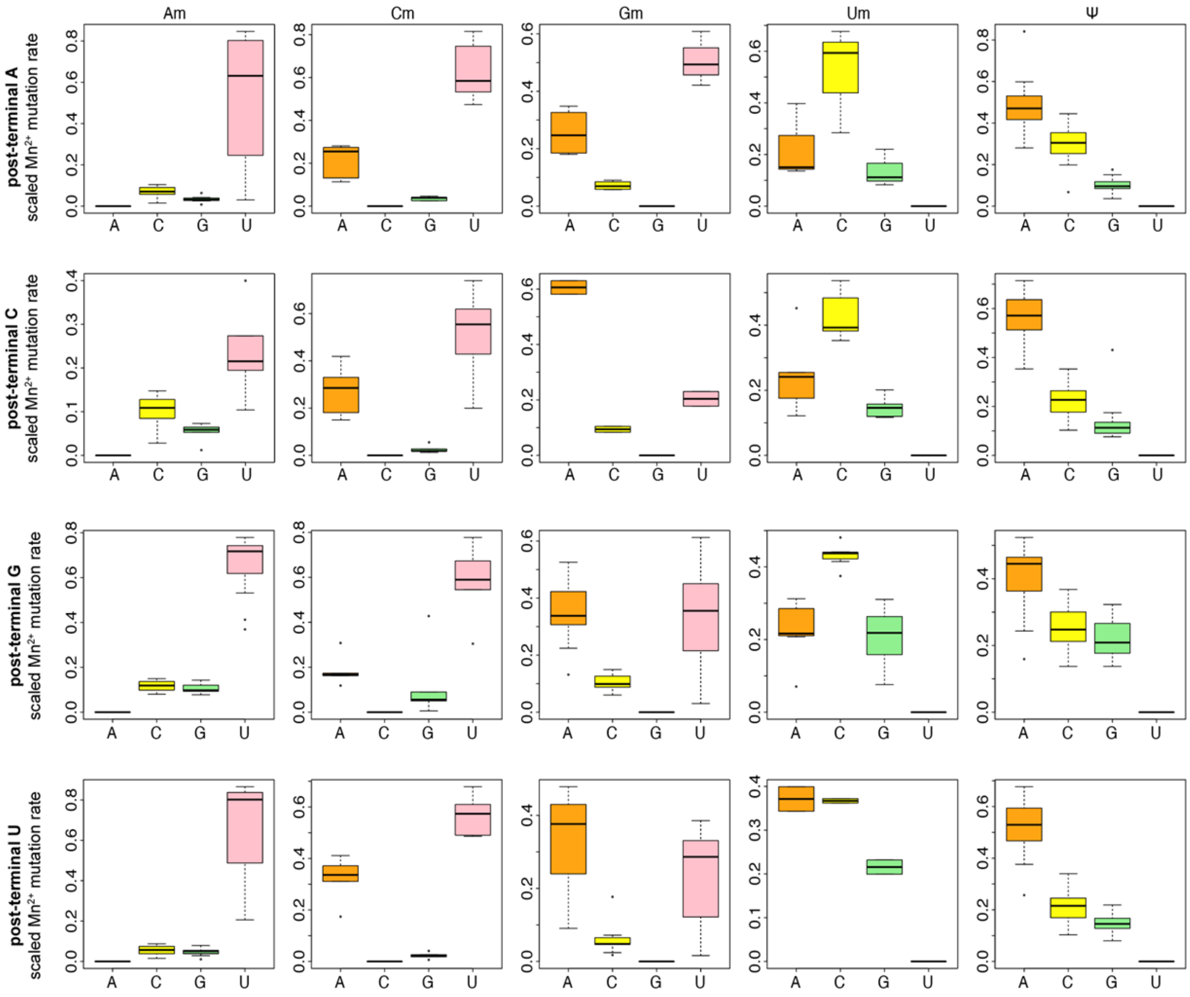


**Figure S8.** Sequence context analysis of manganese (Mn) mutational signatures of Nm and ψ sites in the combined 18S/28S rRNA dataset. The results are sorted by modified nucleotide type (columns) and by the post-terminal nucleotide in the RNA sequence, i.e., the nucleotide immediately 5’ to the position being examined. Each individual plot shows the distribution of marginal (scaled) mutation rates calculated from the Mn sample as a function of the different groups of mutated nucleotides (e.g., the top graph in the Am column shows marginal mutation rates from A to each one of the 4 nucleotides: A to A (no mutation), A to C, A to G, A to U).

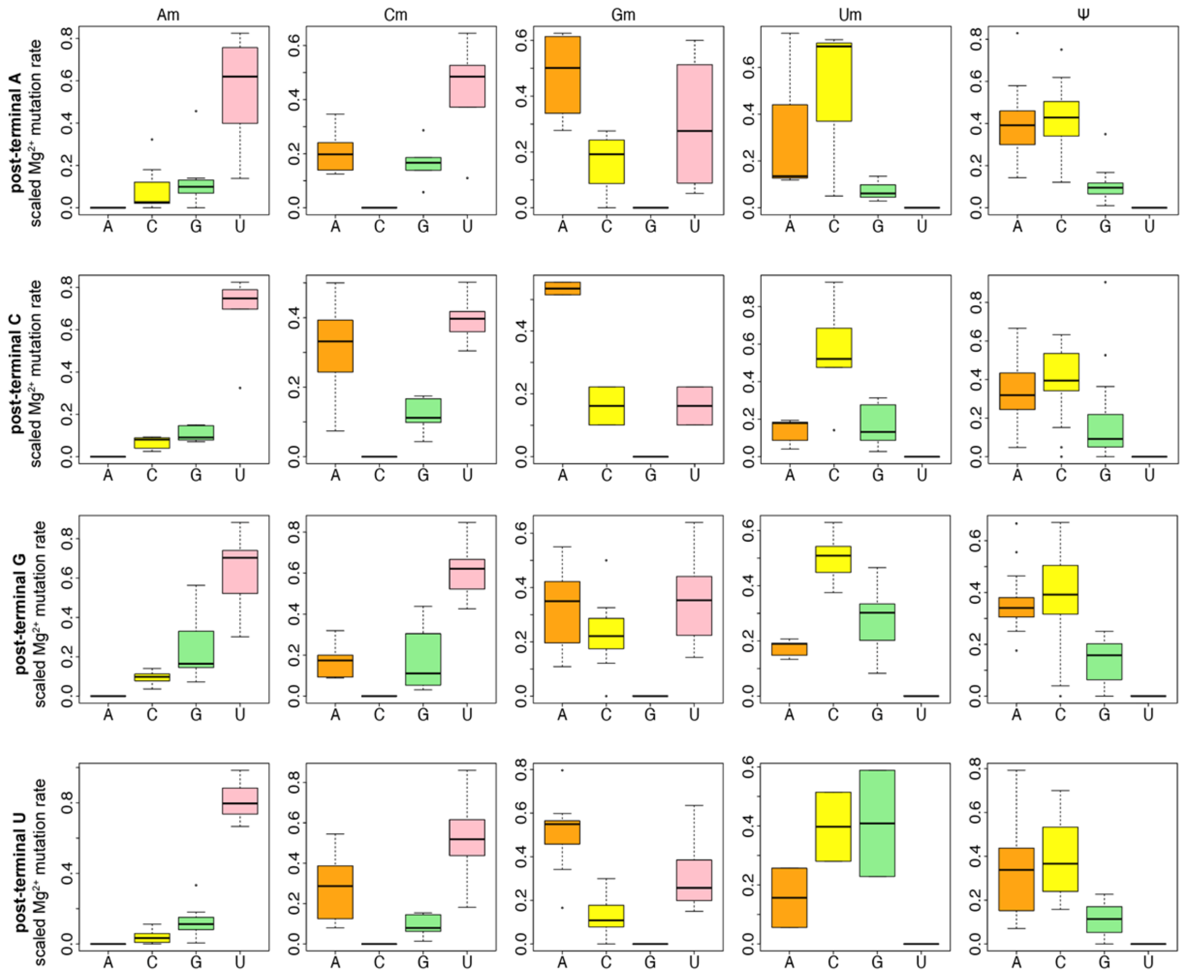


**Figure S9.** Sequence context analysis of magnesium (Mg) mutational signatures of Nm and ψ sites in the combined 18S/28S rRNA dataset. The results are sorted by modified nucleotide type (columns) and by the post-terminal nucleotide in the RNA sequence, i.e., the nucleotide immediately 5' to the position being examined. Each individual plot shows the distribution of marginal (scaled) mutation rates calculated from the Mg sample as a function of the different groups of mutated nucleotides (e.g., the top graph in the Am column shows marginal mutation rates from A to each one of the 4 nucleotides: A to A (no mutation), A to C, A to G, A to U).

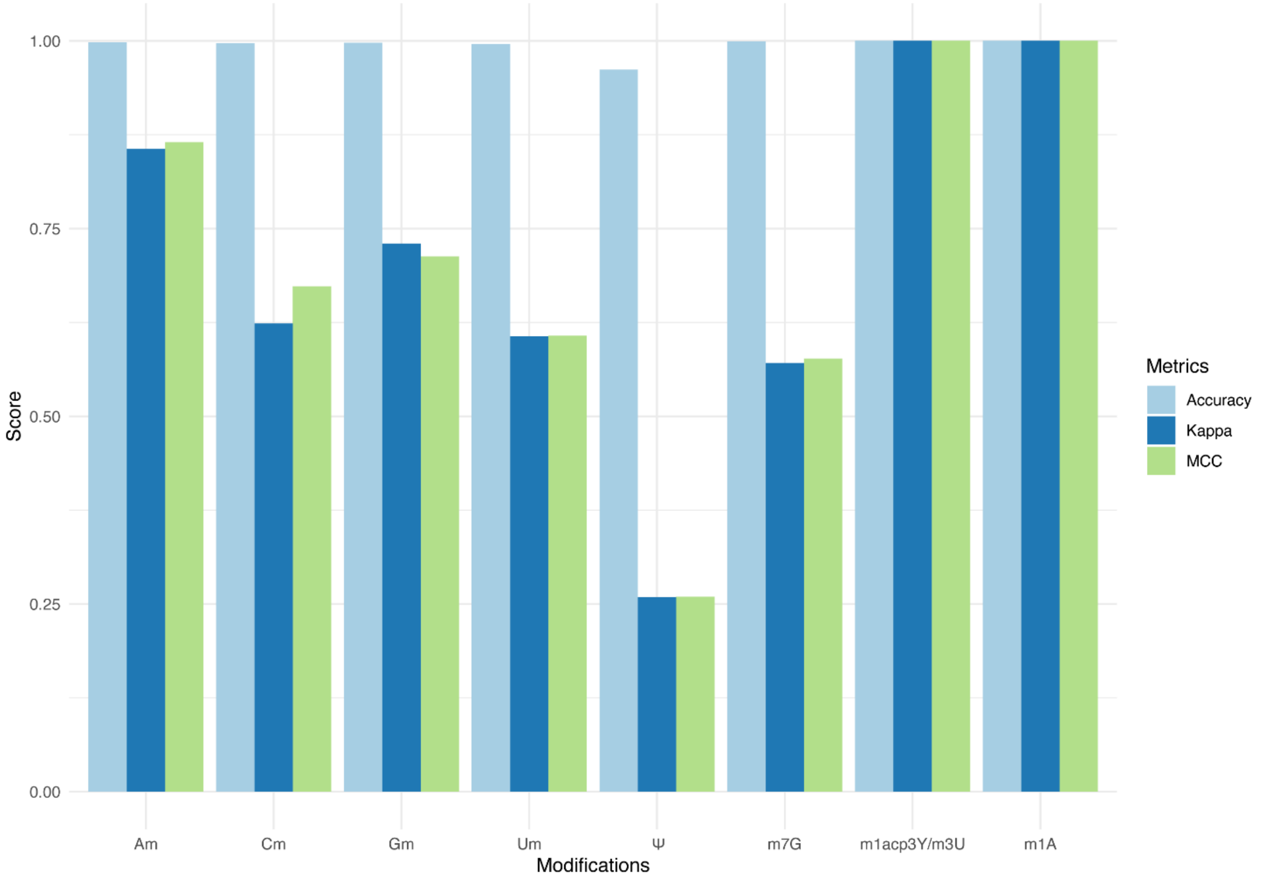


**Figure S10.** Combined benchmarking results of MRT-ModSeq. The bar plot shows the average benchmarking metrics (color legend indicated on the right-hand side for each metric) calculated for each class of RNA modification from the MRT-ModSeq pipeline results over all tested RNAs (human 5.8S rRNA, E coli 16S and 23S rRNAs, S cerevisiae 18S and 25S rRNAs).

**Table S1 – 28S-trained MRT-ModSeq tested on 18S rRNA**

| Mod | Mod sites | Unmod sites | True positives | True negatives | False positives | False negatives | MCC | Kappa | Accuracy |
|---|---|---|---|---|---|---|---|---|---|
| **28S-rRNA Trained Model (Nm & ψ)** | | | | | | | | | |
| **Am** | 13 | 363 | 12 | 363 | 0 | 1 | 0.959 | 0.959 | 0.997 |
| **Cm** | 7 | 448 | 5 | 443 | 5 | 2 | 0.590 | 0.581 | 0.985 |
| **Gm** | 8 | 487 | 4 | 486 | 1 | 4 | 0.628 | 0.611 | 0.990 |
| **Um** | 8 | 338 | 5 | 337 | 1 | 3 | 0.716 | 0.709 | 0.988 |
| **ψ** | 32 | 314 | 13 | 298 | 16 | 19 | 0.371 | 0.371 | 0.899 |

**Table S2 – 18S-trained MRT-ModSeq tested on 28S rRNA**

| Mod | Mod sites | Unmod sites | True positives | True negatives | False positives | False negatives | MCC | Kappa | Accuracy |
|---|---|---|---|---|---|---|---|---|---|
| **18S-rRNA Trained Model (Nm & ψ)** | | | | | | | | | |
| **Am** | 16 | 697 | 15 | 690 | 7 | 1 | 0.794 | 0.784 | 0.989 |
| **Cm** | 15 | 1074 | 10 | 1064 | 10 | 5 | 0.571 | 0.565 | 0.986 |
| **Gm** | 16 | 1294 | 13 | 1276 | 18 | 3 | 0.577 | 0.546 | 0.984 |
| **Um** | 7 | 594 | 6 | 593 | 1 | 1 | 0.856 | 0.856 | 0.997 |
| **ψ** | 55 | 546 | 17 | 512 | 34 | 38 | 0.255 | 0.255 | 0.944 |

### Table S3 – MRT-ModSeq tested on human 5.8S rRNA

| Mod | Mod sites | Unmod sites | True positives | True negatives | False positives | False negatives | MCC | Kappa | Accuracy |
|---|---|---|---|---|---|---|---|---|---|
| **18S+28S-rRNA Trained Model (Nm & ψ)** | | | | | | | | | |
| **Am** | 0 | 31 | 0 | 31 | 0 | 0 | N/A | N/A | 1.0 |
| **Cm** | 0 | 44 | 0 | 44 | 0 | 0 | N/A | N/A | 1.0 |
| **Gm** | 1 | 44 | 1 | 44 | 0 | 0 | 1.0 | 1.0 | 1.0 |
| **Um** | 1 | 35 | 1 | 35 | 0 | 0 | 1.0 | 1.0 | 1.0 |
| **ψ** | 2 | 34 | 0 | 34 | 0 | 2 | N/A | N/A | 0.944 |
| **High-pass filter for base modifications** | | | | | | | | | |
| **m7G** | 0 | 45 | 0 | 45 | 0 | 0 | N/A | N/A | 1.0 |
| **m1acp3Y / m3U** | 0 | 36 | 0 | 36 | 0 | 0 | N/A | N/A | 1.0 |
| **m1A** | 0 | 31 | 0 | 31 | 0 | 0 | N/A | N/A | 1.0 |

### Table S4 – MRT-ModSeq tested on 16S rRNA (*Escherichia coli*)

| Mod | Mod sites | Unmod sites | True positives | True negatives | False positives | False negatives | MCC | Kappa | Accuracy |
|---|---|---|---|---|---|---|---|---|---|
| **18S+28S-rRNA Trained Model (Nm & ψ)** | | | | | | | | | |
| **Am** | 0 | 369 | 0 | 369 | 0 | 0 | N/A | N/A | 1.0 |
| **Cm** | 0 | 339 | 0 | 339 | 0 | 0 | N/A | N/A | 1.0 |
| **Gm** | 0 | 464 | 0 | 464 | 1 | 0 | N/A | 0.0 | 1.0 |
| **Um** | 0 | 303 | 0 | 301 | 2 | 0 | N/A | 0.0 | 0.993 |
| **ψ** | 1 | 302 | 0 | 292 | 10 | 1 | N/A | N/A | 0.964 |
| **High-pass filter for base modifications** | | | | | | | | | |
| **m7G** | 1 | 464 | 1 | 464 | 0 | 0 | 1.0 | 1.0 | 1.0 |
| **m1acp3Y / m3U** | 1 | 302 | 1 | 302 | 0 | 0 | 1.0 | 1.0 | 1.0 |
| **m1A** | 0 | 369 | 0 | 369 | 0 | 0 | N/A | N/A | 1.0 |

### Table S5 – MRT-ModSeq tested on 23S rRNA (*Escherichia coli*)

| Mod | Mod sites | Unmod sites | True positives | True negatives | False positives | False negatives | MCC | Kappa | Accuracy |
|---|---|---|---|---|---|---|---|---|---|
| **18S+28S-rRNA Trained Model (Nm & ψ)** | | | | | | | | | |
| **Am** | 0 | 744 | 0 | 744 | 0 | 0 | N/A | N/A | 1.0 |
| **Cm** | 1 | 621 | 1 | 621 | 0 | 0 | 1.0 | 1.0 | 1.0 |
| **Gm** | 1 | 886 | 1 | 886 | 0 | 0 | 1.0 | 1.0 | 1.0 |
| **Um** | 1 | 576 | 1 | 574 | 2 | 0 | 0.576 | 0.499 | 0.997 |
| **ψ** | 10 | 567 | 3 | 557 | 10 | 7 | 0.248 | 0.246 | 0.971 |
| **High-pass filter for base modifications** | | | | | | | | | |
| **m7G** | 1 | 886 | 0 | 884 | 2 | 1 | -0.001 | -0.001 | 0.997 |
| **m1acp3Y / m3U** | 0 | 577 | 0 | 577 | 0 | 0 | N/A | N/A | 1.0 |
| **m1A** | 0 | 744 | 0 | 744 | 0 | 0 | N/A | N/A | 1.0 |

### Table S6 – MRT-ModSeq tested on Yeast 18S rRNA (*Saccharomyces cerevisiae*)

| Mod | Mod sites | Unmod sites | True positives | True negatives | False positives | False negatives | MCC | Kappa | Accuracy |
|---|---|---|---|---|---|---|---|---|---|
| **18S+28S-rRNA Trained Model (Nm & ψ)** | | | | | | | | | |
| **Am** | 8 | 475 | 7 | 475 | 0 | 1 | 0.93 | 0.93 | 0.998 |
| **Cm** | 3 | 345 | 2 | 345 | 0 | 1 | 0.82 | 0.80 | 0.997 |
| **Gm** | 5 | 454 | 3 | 454 | 0 | 2 | 0.77 | 0.75 | 0.996 |
| **Um** | 2 | 508 | 0 | 508 | 0 | 2 | N/A | 0.00 | 0.996 |
| **ψ** | 14 | 496 | 4 | 486 | 10 | 10 | 0.27 | 0.27 | 0.961 |
| **High-pass filter for base modifications** | | | | | | | | | |
| **m7G** | 1 | 458 | 1 | 458 | 0 | 0 | 1.0 | 1.0 | 1.0 |
| **m1acp3Y / m3U** | 1 | 509 | 1 | 509 | 0 | 0 | 1.0 | 1.0 | 1.0 |
| **m1A** | 0 | 483 | 0 | 483 | 0 | 0 | N/A | N/A | 1.0 |

**Table S7 – MRT-ModSeq tested on 25S rRNA (*Saccharomyces cerevisiae*)**

| Mod | Mod sites | Unmod sites | True positives | True negatives | False positives | False negatives | MCC | Kappa | Accuracy |
|---|---|---|---|---|---|---|---|---|---|
| **18S+28S-rRNA Trained Model (Nm & ψ)** | | | | | | | | | |
| **Am** | 12 | 887 | 8 | 887 | 0 | 4 | 0.81 | 0.80 | 0.996 |
| **Cm** | 7 | 655 | 2 | 655 | 0 | 5 | 0.53 | 0.44 | 0.992 |
| **Gm** | 10 | 956 | 5 | 956 | 0 | 5 | 0.71 | 0.66 | 0.995 |
| **Um** | 8 | 861 | 5 | 861 | 0 | 3 | 0.79 | 0.77 | 0.997 |
| **ψ** | 30 | 839 | 10 | 821 | 18 | 20 | 0.32 | 0.32 | 0.956 |
| **High-pass filter for base modifications** | | | | | | | | | |
| **m7G** | 0 | 966 | 0 | 966 | 0 | 0 | N/A | N/A | 1.0 |
| **m1acp3Y / m3U** | 2 | 867 | 2 | 867 | 0 | 0 | 1.0 | 1.0 | 1.0 |
| **m1A** | 2 | 897 | 2 | 897 | 0 | 0 | 1.0 | 1.0 | 1.0 |

**Table S8 – Benchmarking results for different ML algorithms trained on 18S/28S rRNA uridine (U/Um/ψ) dataset**

| Uridine Original Set | | | | | | | |
|---|---|---|---|---|---|---|---|
| | Naïve Bayes | SVM | Neural Net | kNN | Random Forest | Bagging-J48 | AdaBoost-RT |
| MCC | 0.369 | 0 | 0.418 | 0.383 | 0.511 | 0.547 | 0.511 |
| Kappa | 0.2916 | -0.0039 | 0.4253 | 0.3901 | 0.4818 | 0.5402 | 0.4818 |
| Accuracy | 0.684932 | 0.873288 | 0.876712 | 0.880137 | 0.910959 | 0.914384 | 0.910959 |
| **Uridine Hybridized Set** | | | | | | | |
| | Naïve Bayes | SVM | Neural Net | kNN | Random Forest | Bagging-J48 | AdaBoost-RT |
| MCC | 0.437 | 0.437 | 0.403 | 0.375 | 0.517 | 0.503 | 0.503 |
| Kappa | 0.3979 | 0.3979 | 0.3797 | 0.3518 | 0.5129 | 0.4969 | 0.4969 |
| Accuracy | 0.791096 | 0.791096 | 0.797945 | 0.787671 | 0.873288 | 0.866438 | 0.866438 |
| **Uridine Re-sampled Set** | | | | | | | |
| | Naïve Bayes | SVM | Neural Net | kNN | Random Forest | Bagging-J48 | AdaBoost-RT |
| MCC | 0.276 | 0.460 | 0.441 | 0.407 | 0.502 | 0.527 | 0.436 |
| Kappa | 0.1908 | 0.4171 | 0.4027 | 0.391 | 0.72 | 0.5284 | 0.4458 |
| Accuracy | 0.565068 | 0.797945 | 0.794521 | 0.811644 | 0.883562 | 0.883562 | 0.873288 |
| **Uridine Sub-sampled Set** | | | | | | | |
| | Naïve Bayes | SVM | Neural Net | kNN | Random Forest | Bagging-J48 | AdaBoost-RT |
| MCC | 0.255 | 0.370 | 0.218 | 0.114 | 0.381 | 0.209 | 0.336 |
| Kappa | 0.2114 | 0.2709 | 0.1884 | 0.1204 | 0.3327 | 0.1655 | 0.3029 |
| Accuracy | 0.6130 | 0.6301 | 0.6678 | 0.3630 | 0.7432 | 0.5788 | 0.7432 |

**Table S9 – Benchmarking results for different ML algorithms trained on 18S/28S rRNA adenosine (A/Am) dataset**

| Adenosine Original Set | | | | | | | |
|---|---|---|---|---|---|---|---|
| | Naïve Bayes | SVM | Neural Net | kNN | Random Forest | Bagging-J48 | AdaBoost-RT |
| MCC | 0.539 | -0.010 | 0.897 | 0.947 | 0.947 | 0.897 | 0.952 |
| Kappa | 0.4506 | -0.0056 | 0.8968 | 0.9458 | 0.9458 | 0.897 | 0.9508 |
| Accuracy | 0.9327 | 0.9664 | 0.993884 | 0.9969 | 0.9969 | 0.9938 | 0.9969 |
| **Adenosine Hybridized Set** | | | | | | | |
| | Naïve Bayes | SVM | Neural Net | kNN | Random Forest | Bagging-J48 | AdaBoost-RT |
| MCC | 0.293 | 0.910 | 0.840 | 0.840 | 0.952 | 0.952 | 0.910 |
| Kappa | 0.1578 | 0.906 | 0.272 | 0.8272 | 0.9508 | 0.9508 | 0.906 |
| Accuracy | 0.7615 | 0.9939 | 0.9878 | 0.9878 | 0.9969 | 0.9969 | 0.9939 |
| **Adenosine Re-sampled Set** | | | | | | | |
| | Naïve Bayes | SVM | Neural Net | kNN | Random Forest | Bagging-J48 | AdaBoost-RT |
| MCC | 0.293 | 0.910 | 0.873 | 0.816 | 0.897 | 0.952 | 0.897 |
| Kappa | 0.1578 | 0.906 | 0.8649 | 0.8119 | 0.897 | 0.9508 | 0.897 |
| Accuracy | 0.7615 | 0.9939 | 0.9908 | 0.9878 | 0.9939 | 0.9969 | 0.9939 |
| **Adenosine Sub-sampled Set** | | | | | | | |
| | Naïve Bayes | SVM | Neural Net | kNN | Random Forest | Bagging-J48 | AdaBoost-RT |
| MCC | 0.497 | 0.312 | 0.631 | 0.592 | 0.810 | 0.404 | 0.810 |
| Kappa | 0.3965 | 0.1775 | 0.5697 | 0.5191 | 0.7924 | 0.2804 | 0.7924 |
| Accuracy | 0.9174 | 0.7859 | 0.9572 | 0.9480 | 0.9847 | 0.8685 | 0.9841 |

**Table S10 – Benchmarking results for different ML algorithms trained on 18S/28S rRNA cytidine (C/Cm) dataset**

| Cytidine Original Set | | | | | | | |
|---|---|---|---|---|---|---|---|
| | Naïve Bayes | SVM | Neural Net | kNN | Random Forest | Bagging-J48 | AdaBoost-RT |
| MCC | 0.146 | - | 0.697 | 0.484 | 0.422 | 0.533 | 0.598 |
| Kappa | 0.0867 | 0 | 0.6935 | 0.4616 | 0.3026 | 0.4925 | 0.5274 |
| Accuracy | 0.8190 | 0.9763 | 0.9871 | 0.9806 | 0.9806 | 0.9828 | 0.9849 |
| **Cytidine Hybridized Set** | | | | | | | |
| | Naïve Bayes | SVM | Neural Net | kNN | Random Forest | Bagging-J48 | AdaBoost-RT |
| MCC | 0.131 | 0.262 | 0.397 | 0.374 | 0.562 | 0.573 | 0.632 |
| Kappa | 0.0433 | 0.1285 | 0.2719 | 0.281 | 0.5472 | 0.5587 | 0.6241 |
| Accuracy | 0.7288 | 0.8708 | 0.9446 | 0.9557 | 0.9828 | 0.9741 | 0.9849 |
| **Cytidine Re-sampled Set** | | | | | | | |
| | Naïve Bayes | SVM | Neural Net | kNN | Random Forest | Bagging-J48 | AdaBoost-RT |
| MCC | 0.111 | 0.683 | 0.648 | 0.660 | 0.609 | 0.757 | 0.562 |
| Kappa | 0.0568 | 0.6563 | 0.6323 | 0.6579 | 0.5812 | 0.756 | 0.5472 |
| Accuracy | 0.7608 | 0.9784 | 0.9784 | 0.9828 | 0.9814 | 0.9892 | 0.9828 |
| **Cytidine Sub-sampled Set** | | | | | | | |
| | Naïve Bayes | SVM | Neural Net | kNN | Random Forest | Bagging-J48 | AdaBoost-RT |
| MCC | 0.164 | 0.098 | 0.180 | 0.189 | 0.492 | 0.309 | 0.379 |
| Kappa | 0.0525 | 0.0474 | 0.1201 | 0.1019 | 0.4136 | 0.1745 | 0.2516 |
| Accuracy | 0.5496 | 0.7349 | 0.8599 | 0.7888 | 0.9440 | 0.8211 | 0.8793 |

**Table S11 – Benchmarking results for different ML algorithms trained on 18S/28S rRNA guanosine (G/Gm) dataset**

| Guanosine Original Set | | | | | | | |
|---|---|---|---|---|---|---|---|
| | Naïve Bayes | SVM | Neural Net | kNN | Random Forest | Bagging-J48 | AdaBoost-RT |
| MCC | 0.201 | - | 0.359 | 0.359 | 0.728 | 0.864 | 0.728 |
| Kappa | 0.095 | 0 | 0.3572 | 0.3572 | 0.7245 | 0.8553 | 0.7245 |
| Accuracy | 0.8561 | 0.9889 | 0.9871 | 0.9871 | 0.9945 | 0.9963 | 0.9945 |
| **Guanosine Hybridized Set** | | | | | | | |
| | Naïve Bayes | SVM | Neural Net | kNN | Random Forest | Bagging-J48 | AdaBoost-RT |
| MCC | 0.131 | 0.262 | 0.397 | 0.374 | 0.814 | 0.627 | 0.772 |
| Kappa | 0.0433 | 0.1285 | 0.2719 | 0.281 | 0.7973 | 0.5645 | 0.7465 |
| Accuracy | 0.7288 | 0.8708 | 0.9446 | 0.9557 | 0.9945 | 0.9834 | 0.9926 |
| **Guanosine Re-sampled Set** | | | | | | | |
| | Naïve Bayes | SVM | Neural Net | kNN | Random Forest | Bagging-J48 | AdaBoost-RT |
| MCC | 0.138 | 0.282 | 0.444 | 0.436 | 0.772 | 0.650 | 0.814 |
| Kappa | 0.0478 | 0.1471 | 0.4121 | 0.3593 | 0.7465 | 0.5937 | 0.7973 |
| Accuracy | 0.7472 | 0.8875 | 0.9797 | 0.9686 | 0.9926 | 0.9852 | 0.9945 |
| **Guanosine Sub-sampled Set** | | | | | | | |
| | Naïve Bayes | SVM | Neural Net | kNN | Random Forest | Bagging-J48 | AdaBoost-RT |
| MCC | 0.284 | 0.221 | 0.279 | 0.277 | 0.328 | 0.256 | 0.336 |
| Kappa | 0.1494 | 0.0932 | 0.1448 | 0.1425 | 0.1946 | 0.1231 | 0.2025 |
| Accuracy | 0.8893 | 0.8247 | 0.8856 | 0.8838 | 0.9170 | 0.8653 | 0.9207 |
